# Recent Outbreaks of Shigellosis in California Caused by Two Distinct Populations of *Shigella Sonnei* With Increased Virulence or Fluoroquinolone Resistance

**DOI:** 10.1101/063818

**Authors:** Varvara K. Kozyreva, Guillaume Jospin, Alexander L. Greninger, James P. Watt, Jonathan A. Eisen, Vishnu Chaturvedi

## Abstract

*Shigella sonnei* has caused unusually large outbreaks of shigellosis in California in 2014 – 2015. Preliminary data indicated the involvement of two distinct yet related bacterial populations, one from San Diego and San Joaquin (SD/SJ) and one from the San Francisco (SF) Bay area. Whole genome sequencing of sixty-eight outbreak and archival isolates of *S. sonnei* was performed to investigate the microbiological factors related to these outbreaks. Both SD/SJ and SF populations, as well as almost all of the archival *S. sonnei* isolates belonged to sequence type 152 (ST152). Genome-wide SNP analysis clustered the majority of California (CA) isolates to an earlier described global Lineage III, which has persisted in CA since 1986. Isolates in the SD/SJ population had a novel Shiga-toxin (STX)-encoding lambdoid bacteriophage, most closely related to that found in an *Escherichia coli* O104:H4 strain responsible for a large outbreak. However, the STX genes *(stxla* and *stxlb)* from this novel phage had sequences most similar to the phages from *S. flexneri* and *S. dysenteriae*. The isolates in the SF population yielded evidence of fluoroquinolone resistance acquired via the accumulation of point mutations in *gyrA* and *parC* genes. Thus, the CA *S. sonnei* lineage continues to evolve by the acquisition of increased virulence and antibiotic resistance, and enhanced monitoring is advocated for its early detection in future outbreaks.

## IMPORTANCE

Shigellosis is an acute diarrheal disease causing nearly a half-million infections, 6,000 hospitalizations, and 70 deaths every year in the United States. *Shigella sonnei* caused two unusually large outbreaks of shigellosis in 2014 – 2015 in California. We applied whole genome sequence analyses to understand the pathogenic potential of bacteria involved in these outbreaks. Our results suggest that a local *S. sonnei* clone has persisted in California since 1986. Recently, a derivative of the original clone acquired the ability to produce Shiga-toxin via exchanges of bacteriophages with other bacteria. Shiga toxin production is connected with more severe disease including bloody diarrhea. A second derivative of the original clone recovered from the San Francisco Bay area showed evidence of gradual acquisition of antimicrobial resistance. These evolutionary changes in *S. sonnei* populations must be monitored to explore the future risks of spread of increasingly virulent and resistant clones.

## Introduction

Shigellosis is an acute gastrointestinal infection caused by bacteria belonging to the genus *Shigella*. Shigellosis is the third most common enteric bacterial infection in the United States with 500,000 infections, 6,000 hospitalizations, and 70 deaths each year [1]. There are four *Shigella* species which cause shigellosis: *S. dysenteriae, S. flexneri, S. boydii*, and *S. sonnei* [2]. *S. dysenteriae* is considered to be the most virulent species especially *S. dysenteriae* type 1 serotype due to its ability to produce a potent cytotoxin called Shiga-toxin. *S. flexneri, S. boydii*, and *S. sonnei* generally do not produce Shiga-toxin (STX) and, therefore, cause mild forms of shigellosis [2, 3]. Shiga-toxins can also be produced by Shiga toxin-producing *Escherichia coli* (STEC). Two types of STX are known in STEC: STX 1 which differs by a single amino acid from STX in *S. dysenteriae* type 1, and STX 2 which shares only about 55% amino-acid similarity with STX 1 [4]. STX-operons for both type 1 and 2 STX consist of the *stxA* and *stxB* subunit genes, which encode the AB_5_ holotoxin [5]. Rarely, Shiga-toxin genes *stx* can be transferred to non-STX-producing *S. flexneri* and *S. sonnei* by means of lambdoid bacteriophage either from STEC or *S. dysenteriae* [3, 6, 7], providing those strains with the ability to cause more severe disease. Infections caused by bacterial species that produce STX often lead to hemorrhagic colitis and may cause serious complications, like hemolytic uremic syndrome (HUS) [8].

*S. sonnei* has caused two large outbreaks in California (CA) in 2014-2015. The first outbreak with distinct clusters in the San Diego and San Joaquin (SD/SJ) areas was caused by STX-producing isolates [9]. Another CA *S. sonnei* outbreak occurred within the same time frame as the two clusters described above, but was caused by a STX-negative *S. sonnei* strain and was confined primarily to the San Francisco Bay (SF) area [10]. We performed whole genome sequencing of sixty-eight outbreak and archival isolates to gain further insights into the microbiological factors associated with these shigellosis outbreaks. We examined the phylogeny of local *S. sonnei* isolates and also performed comparison of CA isolates with global *S. sonnei* clones. The results provided new insight into likely origin of current CA *S. sonnei* isolates and evolutionary changes to acquire more virulence and antibiotic resistance.

## RESULTS

### *In silico* MLST and hq SNP Analysis

Sequences of 7 house-keeping genes were extracted from genomic sequences of all isolates and their allele numbers were identified using the CGE MLST database [11]. The sequence type (ST) was assigned to each isolate based on combination of identified alleles. All outbreak and archival CA *S. sonnei* isolates had identical sequences for 7 MLST loci and were assigned to the same sequence type 152 (ST152), with the exception of one historical isolate C97 identified as ST1502. ST1502 differs from ST152 by a single allele. *S. sonnei* ST 152 was previously found in Germany in 1997 and China in 2009 [12]. In the MLST database, there is also a record of ST152 isolated in the United Kingdom in 2012, though it is not clear if this ST was assigned to *E. coli* or *Shigella* species.

Based on genome-wide SNP analysis, isolates from CA clustered into several clades (Figure 1A). All STX-positive isolates from two recent outbreak-related San Diego (SD) and San Joaquin (SJ) epidemiological groups clustered together and differed from each other by 0-11 SNPs. The member of SD epidemiological group C24 had 0 SNP differences with several SJ isolates (C101, C114-C119, C122, C123, C125, C128, C129), indicating direct relatedness of the isolates from the SD and SJ epidemiological clusters. The only STX-negative isolate C130 from SJ had 48 SNPs differences from the closest member of STX-positive SD/SJ group.

**Figure 1.**
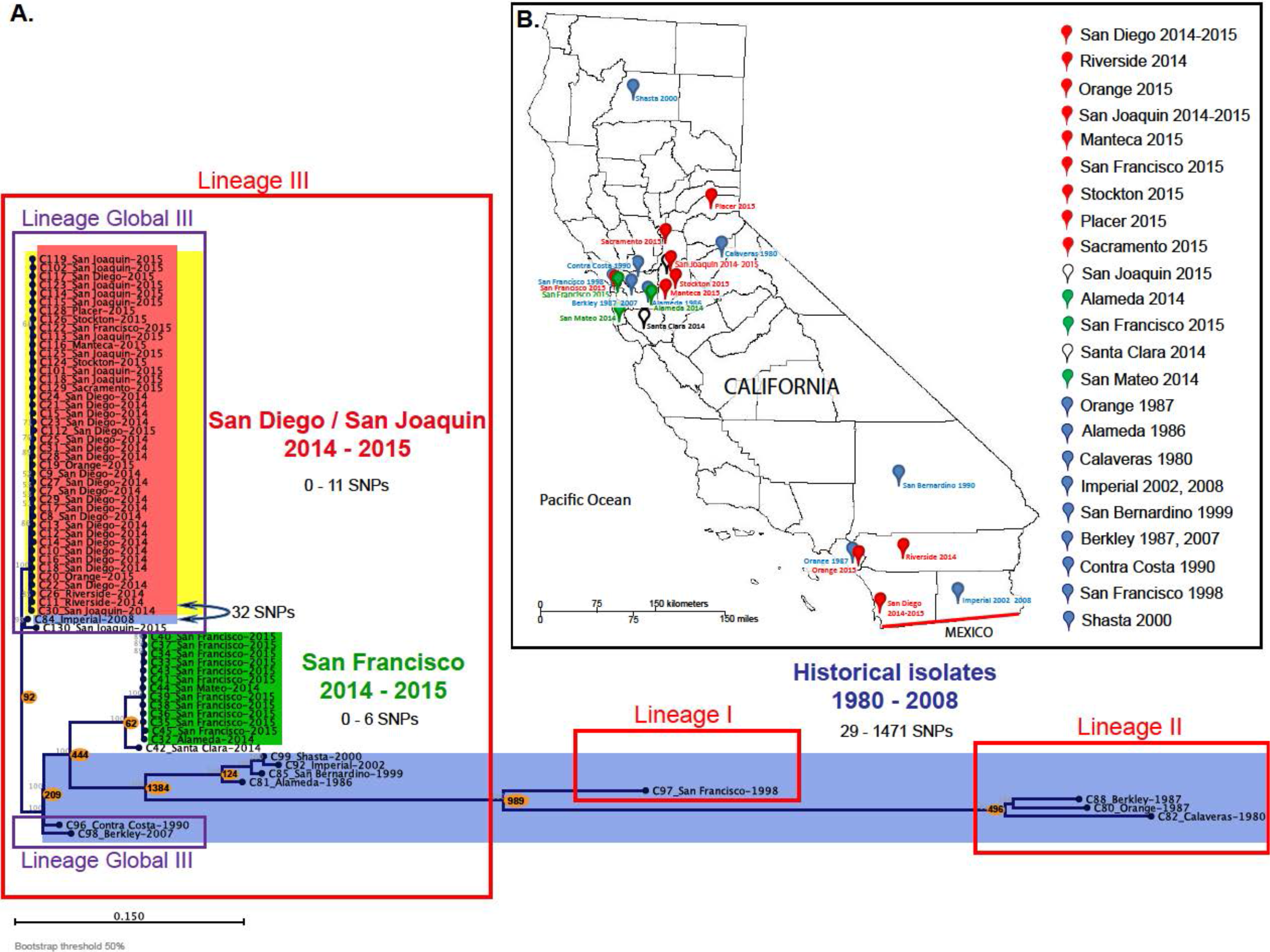
Local and global phylogeny of CA *S. sonnei*. A. Maximum-likelihood phylogenetic tree based on genome-wide hqSNP differences between STX1-producing *S. sonnei* from San Diego and San Joaquin outbreak, STX-negative *S. sonnei* from San Francisco outbreak and historical isolates from CA. Background color: Red-San Diego/San Joaquin (SD/SJ) outbreak; Green-San Francisco outbreak; Blue-historical isolates; Yellow-STX-positive isolates. Frame line color: Red-designates major lineages I-III by Holt et al.; Purple-sub-lineage Global III by Holt et al. Node labels are displayed as “Isolate ID_Geographic location in California-Date of isolation”. Numbers in orange circles correspond to the numbers of SNPs different between the closest isolates of the groups. B. Geographical distribution of the CA *S. sonnei* isolates. The international boundary between the US and Mexico is drawn in red. Color of the pins corresponds to the following groups of isolates: red-SD/SJ population, green-SF population, blue-historical isolates, white-modern sporadic isolates

Isolates from the contemporary outbreak in San Francisco (SF) formed a separate cluster (Figure 1A), and had more SNP differences with the SD/SJ cluster (251 SNPs) than with the historical isolate C96 from Contra Costa County, isolated in 1990 (219 SNPs). Isolate C42 from Santa Clara County (2014) was assigned to the SF outbreak based on PFGE (Table S1), but differed by 62 SNPs from the closest SF outbreak member. C42 clustered closer to the SF outbreak than to any other contemporary or historical isolate, therefore was considered to be a part of the SF population.

The majority of historical isolates were genetically distinct from recent isolates except C84 from Imperial County (2008), which differed by 32 SNPs from closest member of SD/SJ STX-positive cluster and by 28 SNPs from STX-negative SJ isolate C130. This is surprisingly small number of SNPs, when compared with the distance between the SD/SJ cluster and other historical isolates (100-1474 SNPs) or even to the contemporary SF outbreak cluster (251 SNPs).

### Global phylogeny of CA S. sonnei

Genomes of CA *S. sonnei* were compared with global *S. sonnei* clones reported by Holt et al. (Table S2) [13] by genome-wide SNP analysis, which is expected to provide better resolution of lineages than MLST. The clustering of global strains in our analysis showed the same general patterns as in the original publication (Fig. S1). Both modern *S. sonnei* populations and the majority of the historical isolates (1986-2008) clustered with Lineage III strains. The European isolate from the UK belonging to the Global III Lineage isolated in 2008 (the most recent from the analyzed collection by Holt et al.), was the closest global strain to both the SD/SJ and the SF *S. sonnei* populations. Historical isolates C96, C98, and C84 (from 1990, 2007, and 2008, correspondingly), also clustered together with the Global III Lineage. Another clade within Lineage III was formed by an isolate from Mexico (1998) and a group of historical CA *S. sonnei* isolates obtained between 1986 and 2002. Three CA isolates from 1980-1987 clustered together with Lineage II. A single historical isolate C97, which also had a unique MLST pattern ST1502, was grouped with Lineage I strains. No Lineage IV isolates were found in CA (Figure 1A, Fig. S1)

### COG and pfam clustering of CA *S. sonnei* genomes

Hierarchical clustering based on COG (the Clusters of Orthologous Groups of proteins) and pfam (protein domains) profiles was used as an alternative method for examining similarity and relatedness of strains. This protein family profile clustering was carried out with representative genomes of CA *S. sonnei* isolates (C7, 113, C123-SD/SJ population; C39- SF population; C84, C88, C97- historical isolates) and other publicly available genomes of *Shigella* species and different *E. coli* serotypes. Two different clustering methods were used here – one based on presence/absence and one based on abundance (in both cases of COG and pfam groups). The COG-based approach showed all CA *S. sonnei* isolates clustering together with *E. coli* and separate from other *S. sonnei* isolates, with the exception of *S. sonnei* strain 1DT-1, which also clustered with *E. coli* (Fig. S2A). CA *S. sonnei* clustered closer with non-O157 serotypes of *E. coli*, particularly with O104:H4 STEC strain (which caused a large outbreak in Germany and other European countries in 2011) than with *E. coli* O157:H7. Similar results were seen in the hierarchical clustering based on Pfam profiles (Fig. S2B). Except in the pfam clustering two CA *S. sonnei* isolates (one of the historical isolates and one San Francisco isolate) clustered with other *S. sonnei* from the database, while all other CA isolates clustered with *E. coli*, as previously.

To determine whether specific parts of genome were shared between CA *S. sonnei* and *E. coli* and contributed to CA *S. sonnei* clustering with *E. coli*, we searched for the genes in the genome of CA *S. sonnei* which had homologs in *E. coli* O104:H4 strains, but not in other *S. sonnei* from the database. The genes which were found to be shared between *E. coli* O104:H4 and STX-positive CA *S. sonnei* isolates (representative, C7 and C123) belonged to either the STX-bacteriophage characterized here (for both C7 and C123), or to a *bla*_TEM-1_-encoding plasmid (in case of C123) (Table S3). The CA *S. sonnei* isolates lacking STX-bacteriophage didn't have any genes which were shared with *E. coli* O104:H4, but not with other *S. sonnei*.

We also performed a recombination test on alignment of genome-wide hqSNPs using the Recombination Detection Program v. 4 (RDP4) [14]. None of the used recombination detection algorithms included in the software package detected recombination events between CA *S. sonnei* and *E. coli* O104:H4. Thus, no other gene exchange events, except for STX-bacteriophage transfer, were detected between *E. coli* O104:H4 and CA *S. sonnei*.

A phylogenetic tree of genomes was constructed from a concatenation of a set of 37 core phylogenetic marker genes using Phylosift (Fig. S3A) [15]. In this tree, all *S. sonnei* including CA isolates grouped together, and separate from *E. coli* with one exception. The exception was *S. sonnei* strain 1DT-1, which grouped in a clade with *E. coli*. Both nucleotide (Fig. S3A) and amino acid (AA) sequence (data not shown) trees generated with Phylosift showed the same result. Whole genome clustering based on genome-wide SNPs in both coding and non-coding areas of the genome (Fig. S3B) supported the results of Phylosift: all *S. sonnei* isolates, including ones from CA, clustered together, except for *S. sonnei* 1DT-1.

The difference between protein family profile clustering and concatenated marker gene phylogeny is striking. In the protein family profile clustering CA *S. sonnei* isolates group with *E. coli* O104:H4 than with other *S. sonnei* but in the concatenated marker gene phylogeny CA *S. sonnei* and *E. coli* do not group together. At first, to explain this contrast we considered two plausible biological explanations:1) convergent gene gain or loss could have led to high similarity in terms of protein family profiles between CA *S. sonnei* and *E. coli* even though they are not closely related, or 2) considering the hypothesis of origination of all *Shigella* species from *E. coli* [16], CA *S. sonnei* could be shearing higher similarity of protein family profiles with *E. coli* because it represents a more archaic lineage of *S. sonnei* species, which haven't diverged from *E. coli* as much as the other *Shigella* strains found in the NCBI and IMG databases did.

However, neither of these explained all the results from these analyses. Closer examination of some of the results led us to consider a third possibility – the observed contrast was an artifact of analysis. Upon further review, we found that CA isolates and *S. sonnei* strain 1DT-1 similar to *E. coli* O104:H4 had low gene count for several Transposases (COG2963, COG2801, COG3547, COG3335, COG4584, COG3385) and a DNA replication protein DnaC (COG1484), while other *Shigella* spp. isolates in the database had those genes in abundance (Fig. S4). This indicated that a few transposable elements skewed the clustering of CA *S. sonnei* towards *E. coli*. When only the presence/absence of the COG and pfam elements was taken into the account, COG-and pfam-based clustering appeared to be congruent with nucleotide phylogeny: CA *S. sonnei* have clustered with other *S. sonnei* strains in the database (Figure 2; Fig. S5). Therefore, we would like to emphasize that genome clustering based on gene presence/absence as well as abundance is prone to the issues described above and shouldn't be used to infer phylogenies.

**Figure 2.**
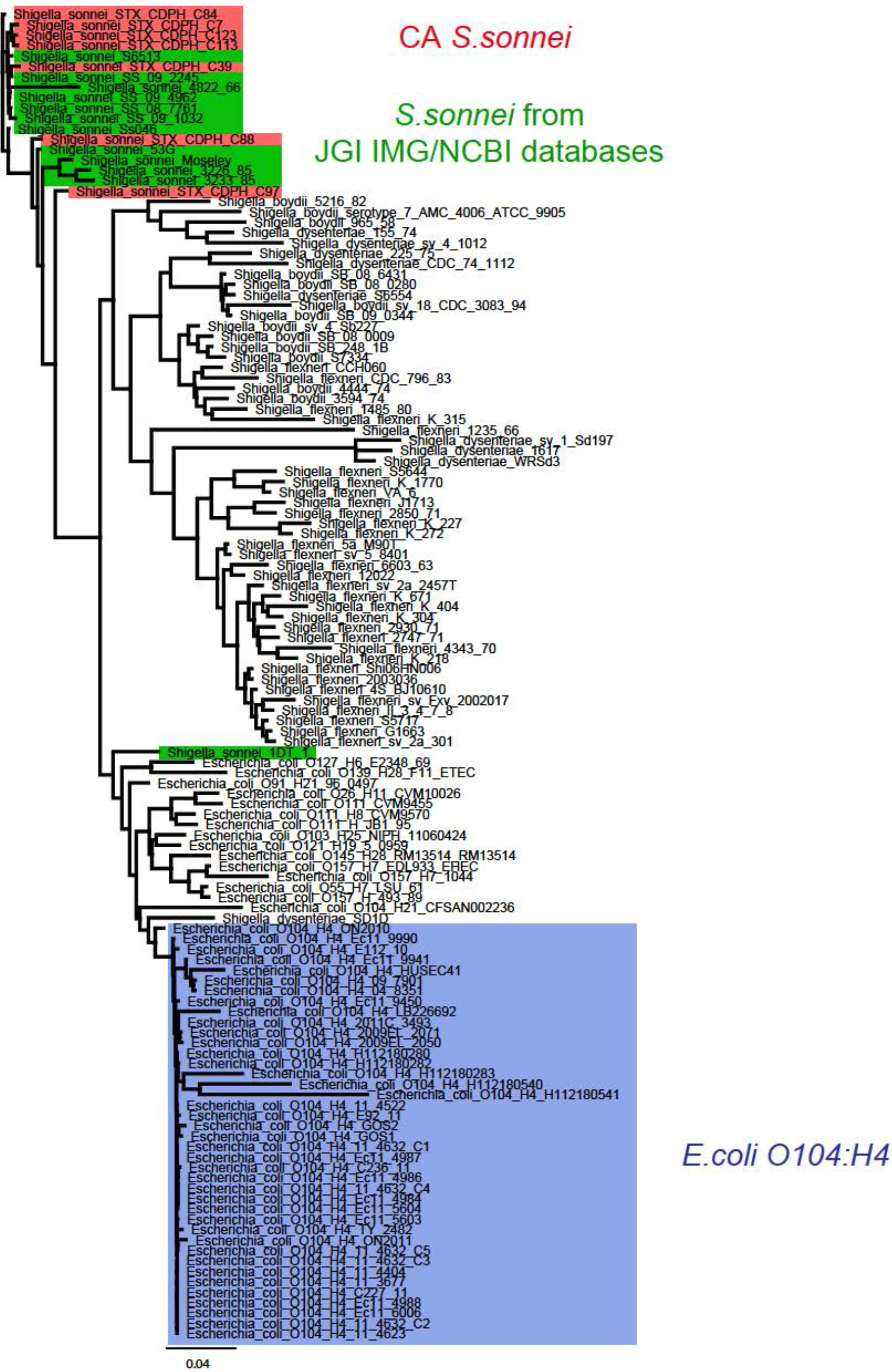
Comparison of CA *S. sonnei* with *E. coli* strains and other publicly-available genomes of *S. sonnei*. Revised Maximum Likelihood clustering of CA representative isolates with *E. coli* and other *Shigella* species from IMG JGI database based on COG profiles (presence/absence). Background color: Red-representative *S. sonnei* from California; Green-other *S. sonnei* from JGI IMG database; Blue-*E. coli* O104:H4 strains from JGI IMG database.

### STX-encoding bacteriophage from CA *S. sonnei*

All isolates from SD/SJ outbreak possessed STX-1 subunits A and B genes (*stxlA, stxlB*) (Figure 3). None of the historical isolates or recent isolates related to the San Francisco outbreak had *stx* genes. The STX in the SD/SJ population was encoded by a novel lambdoid bacteriophage. This 62.8kb bacteriophage had identical sequence and was integrated into a chromosomal *wrbA* site in all STX-positive isolates. A modern STX-negative *S. sonnei* isolate C130 from SJ had an intact *wrbA* site; therefore, we conclude that it represents an ancestral STX-negative lineage for the SD/SJ population.

**Figure 3.**
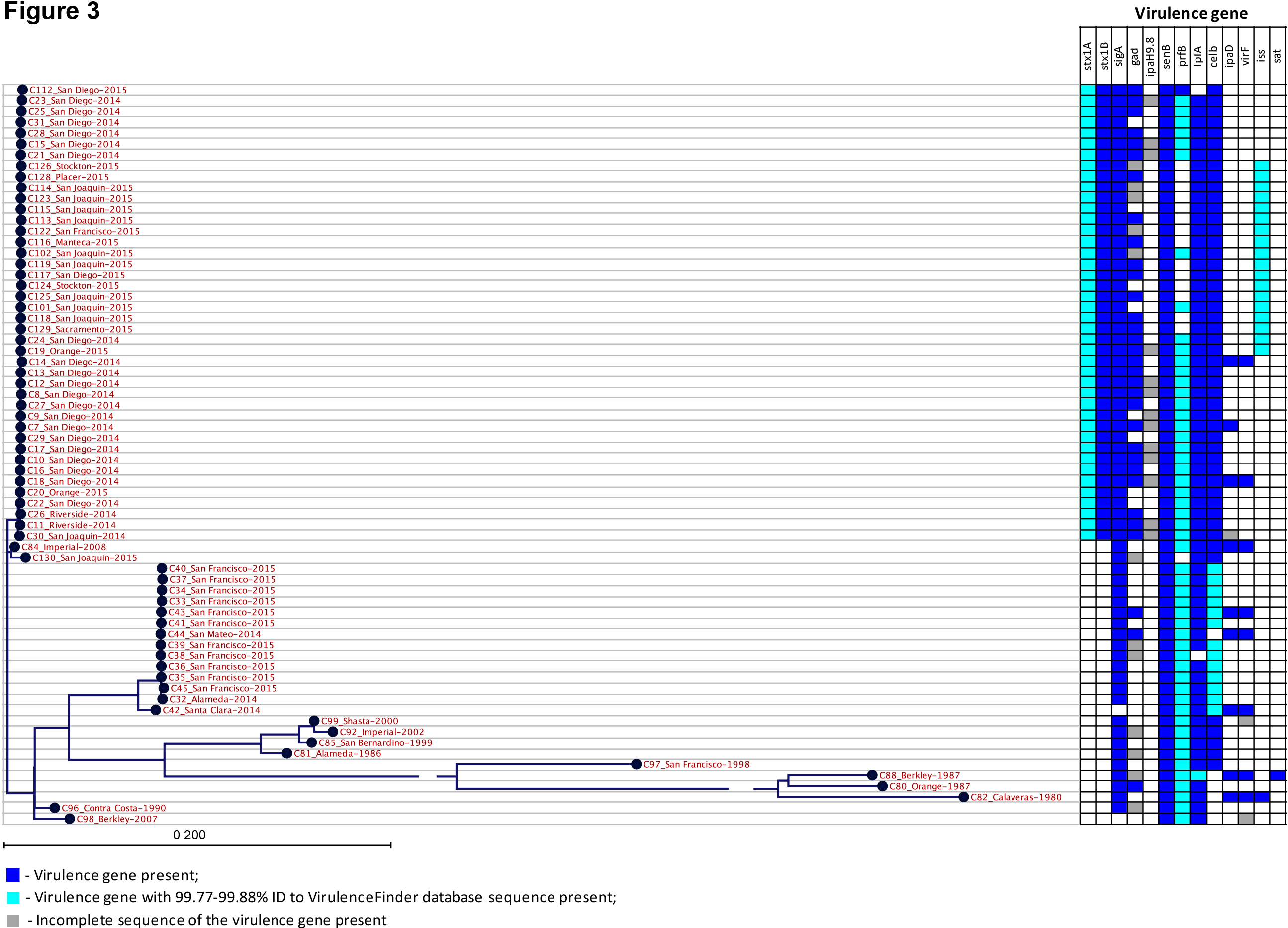
Virulence determinants distribution in California *S. sonnei*. Virulence determinants names: *stx1A*- Shiga toxin 1, subunit A, variant a; *stx1B*- Shiga toxin 1, subunit B, variant a; *sigA*- Shigella IgA-like protease homologue; *gad*- Glutamate decarboxylase; *ipaH9.8*- Invasion plasmid antigen; *senB*- Plasmid-encoded enterotoxin; *prfB*- P-related fimbriae regulatory gene; *lpfA*- Long polar fimbriae; *celb*- Endonuclease colicin E2; *ipaD*- Invasion protein *S. flexneri*; *virF*- VirF transcriptional activator; *iss*- Increased serum survival; *sat*- Secreted autotransporter toxin.

In order to better understand the history of the novel STX-phage in the SD/SJ *S. sonnei* population, we carried out a series of comparative analyses. First, the Phage Search Tool (PHAST) revealed that the best match for the novel CA STX1-phage was the STX2-encoding P13374 phage found in O104:H4 STEC, an outbreak strain that caused a large outbreak in Germany and other European countries in 2011 [17]. This finding was supported by Blast searcher which revealed that the bacteriophage from the CA isolates was very similar to two phages found in strains from the European *E. coli* O104:H4 outbreak [P13374 phage (NC_018846)] with 97% identity for 88% of the query search coverage and the phage in *E. coli* strain 2011C-3493 (CP003289.1) with 97% identity for 93% of the query coverage. CA STX1-phage also showed very high similarity with stx2-encoding phages from *E. coli* O104:H4 strains 2009EL-2050 (CP003297.1) and 2009EL-2071 (CP003301.1) from Georgia, 2009 [17] with 98% shared identity at 92% query length. A recently characterized STX1-encoding *S. sonnei* phage 75/02_Stx from Hungary (KF766125.2) was also shown to share large co-linear regions with STX2-prophages from *E. coli* O104:H4 and O157:H7 [5]. However, the Hungarian phage only partially resembled CA STX1-phage with 99% of shared identity at 62% query coverage.

In order to better understand the phylogenetic relatedness of CA STX1-phage to other phages, we generated phylogenetic trees of integrase genes. This revealed that the CA STX1-phage grouped evolutionarily with other STX-encoding *E. coli* phages, particularly with phages from *E. coli* O104:H4, but not with ones from *Shigella* species (Fig. S6). Figure S6 shows the phylogeny based on Integrase amino-acid sequences; similar results were seen with nucleotide based trees (not shown). This integrase phylogenetic analysis confirmed the relatedness of the CA STX-phage to the phage found in pandemic *E. coli* O104:H4 from Germany, 2011. It also showed that bacteriophages with integrase genes related to phage from European outbreak *E. coli* O104:H4 were also found earlier in Europe (Norway, 2007 and Georgia, 2009).

We also used progressive Mauve to make and analyze whole phage alignments. This revealed that the STX1-phage in CA isolates more closely resembled STX-coding phages in *E. coli* than phages in *S. sonnei* or *S. flexneri* (Figure 4A, 4B). The stx2-encoding phage from *E. coli* O104:H4 strain 2011C-3493 was the closest to CA STX-phage according to Neighbor-Joining tree built based on the estimate of the shared gene content from progressiveMauve alignment. Notably, *E. coli* strain 2011C-3493 was isolated in the US from a patient with the history of travel to Germany and shown to be a part of German outbreak [18].

**Figure 4.**
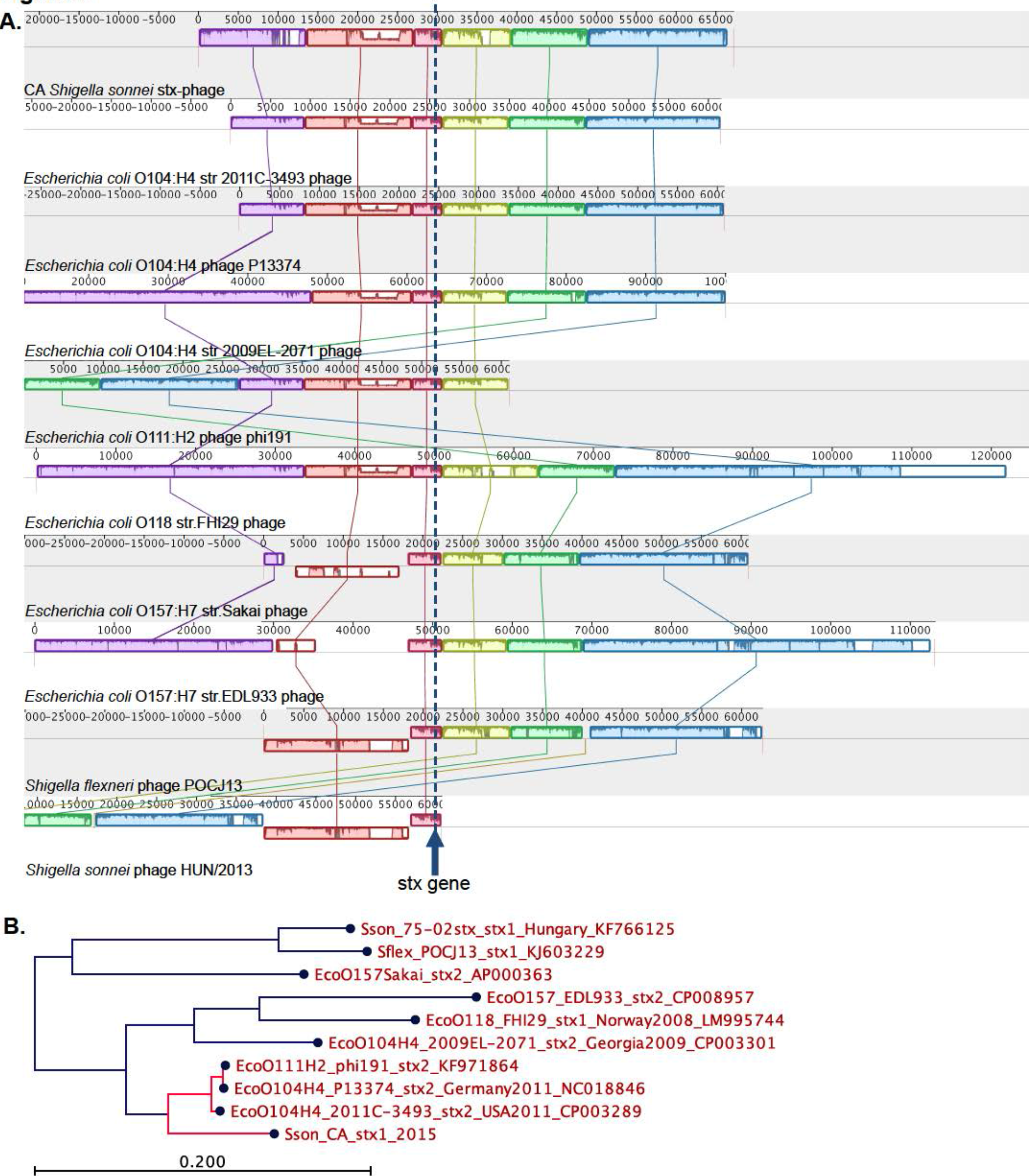
Phylogeny of Shiga-toxin-encoding phage from SD/SJ *S. sonnei*. A. progressiveMauve alignment of phage sequences. Sequences are centered by the *stxA* gene. B. A neighbor-joining tree based on an estimate of the shared gene content from the progressiveMauve alignment.

Examination of the distribution of CA STX1-bacteriophage-related genes in different *E. coli* and *Shigella* serovars in the IMG database showed that the CA STX-phage genes were also present in *E. coli* O104:H4 isolates, the majority of which were directly related to the German outbreak. Strain ON2010 of *E. coli* O104:H4 from Canada (2010), which was not linked to the German outbreak [19], showed no presence of CA STX-phage genes (Fig. S7).

In several CA isolates we identified the bacteriophages known to be associated with STX, but which were missing actual *stx* genes. According to PHAST output, the isolate C7 possessed an intact STX2-converting phage 1717 (NC_011357), isolate C113 had an incomplete STX2-converting phage 86 (NC_008464), and isolates C16 harbored a questionable STX2-converting phage I (NC_003525), while none of them contained the *stx* gene sequence (Fig. S8). We propose that the *stx* genes have been gained and lost during the evolution of *S.sonnei* in California.

### STX-holotoxin and other virulence determinants CA *S.sonnei*

Even though the phage from German outbreak STEC strain was identified as the most closely related to the CA STX1-phage, it carries the *stx2* gene, while CA phage encodes the STX1 toxin. The STX-operon in CA isolates (holotoxin genes *stx1a* and *stx1b*) was identical to STX operons in *S. flexneri* phage POCJ13 (KJ603229.1) and several *S. dysenteriae* strains, including *S. dysenteriae* type 1 (M19437.1), but differed from STX operon in the majority of *E. coli* strains. The CA *S. sonnei* STX-operon had 1 SNP difference with other STX1-encoding *S. sonnei* bacteriophages (Accession ## KF766125.2 and AJ132761.1). Similarity of different regions of CA STX1-phage to various phages can be explained by the mosaic structure of STX lambdoid phages due to frequent recombinations and modular exchange with other lambdoid phages [20]. Moreover, there is evidence that it could be quite common for lambdoid phages of *S. sonnei* to have an STX genes from *S. dysenteriae* and the rest of the phage genes resembling STEC or EHEC *E. coli* strains [3, 5].

Besides STX, the following virulence genes were detected in *S. sonnei* from CA: Shigella IgA-like protease homologue (*sigA*), glutamate decarboxylase (*gad*), plasmid-encoded enterotoxin (*senB*), P-related fimbriae regulatory gene (*prfB*), long polar fimbriae (*lpfA*), endonuclease colicin E2 (*celb*), invasion protein *S. flexneri* (*ipaD*), VirF transcriptional activator (*virF*), and increased serum survival gene (*iss*). The distribution of these virulence genes in different *S.sonnei* populations from CA is presented in Figure 3. The increased serum survival virulence gene *iss* was mainly found in more recent of the SD/SJ population of isolates, and in a single historical isolate from 1980, however the *iss* alleles in modern and historical isolates were different.

### Antimicrobial resistance of CA *S.sonnei*

Acquired antibiotic resistance (ABR) markers to the following classes of antimicrobials were identified: beta-lactams (genes *bla*>_TEM-1_; *bla*_OXA-2_), aminoglycosides (*aph(3′′)-Ib*; *aph(6)-Id*; *aac(3)-IId*; *aadA1*; *aadA2*), macrolides (*mphA*), sulphonamides (*sul1*; *sul2*), phenicols (*catA1*), trimethoprim (*dfrA1; dfrA8; dfrA12*), and tetracycline (*tetA*; *tetB*) (Figure 5). The majority of the isolates possessed genes predicted to provide resistance to aminoglycosides, sulphonamides, trimethoprim, and tetracycline. In addition, a phenotypic resistance to penicillins mediated by TEM-1 β-lactamase gene occurred in 16 isolates. The correlation between the genotype and susceptibility phenotype is presented in the Table S4. In 100% of cases, the presence of antibiotic resistance determinants correlated with the expected non-susceptible phenotype. The 92.9% of the isolates lacking the resistance determinant were susceptible to corresponding antimicrobials. A variant of the gene of aminoglycoside-modifying enzyme *aac(3)-IId* was found in a single *S. sonnei* isolate C98 from CA; the *aac(3)-IId*-like gene in isolate C98 shared 99.8% identity with previously described variant of this gene.Acetyltransferase *aac(3)-IId* was shown previously to confer resistance to gentamicin and tobramycin in *E. coli* isolates [21], while a *S. sonnei* isolate harboring a variant of this gene was intermediate to tobramycin and resistant to gentamicin.

**Figure 5.**
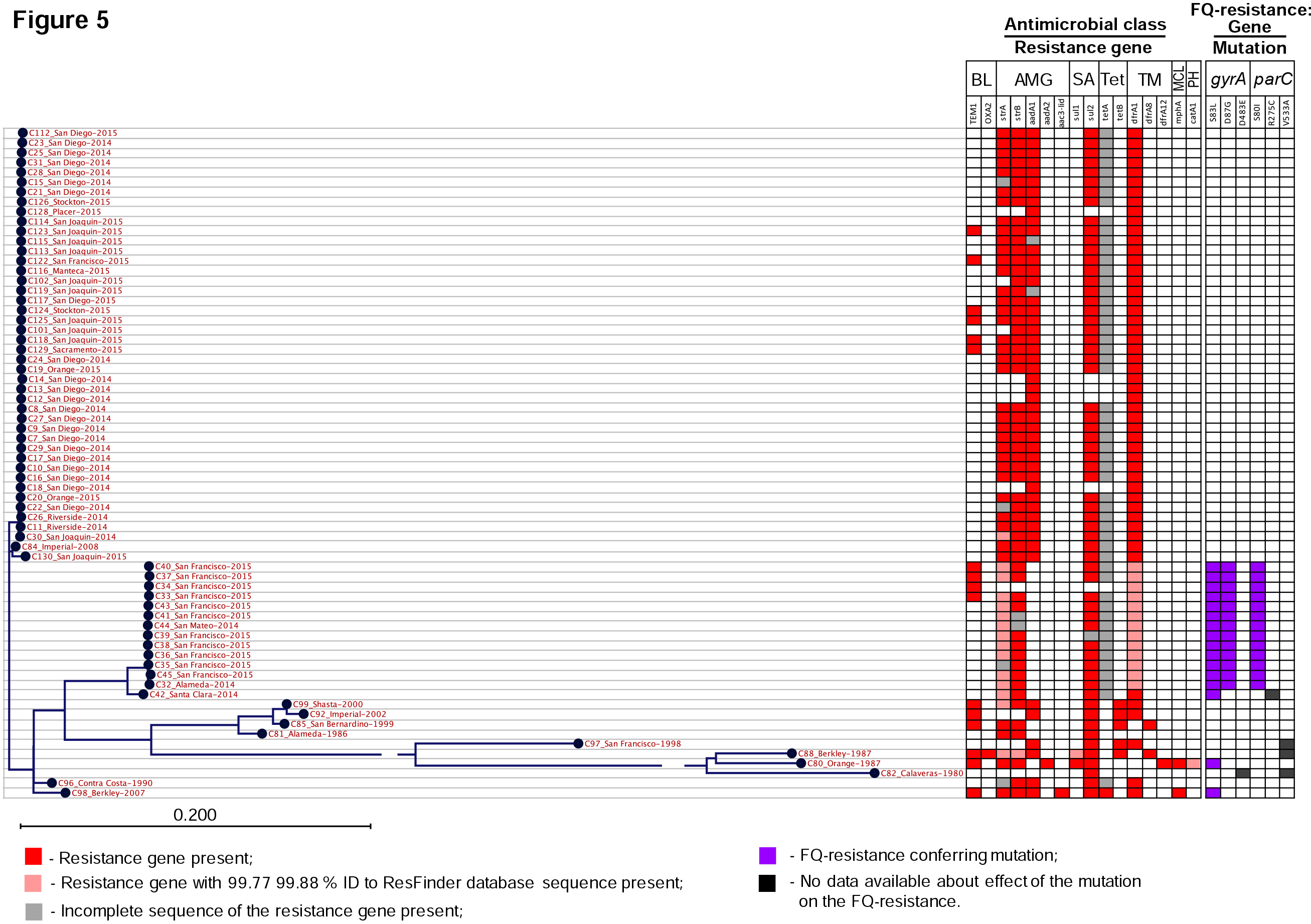
Antibiotic resistance determinants distribution in CA *S. sonnei*. Abbreviations of antimicrobial classes: BL-Beta-lactams; AMG-Aminoglycosides; SA-Sulphonamides; Tet-Tetracycline; TM-Trimethoprim; MCL-Macrolide; PH-Phenicols; FQ-fluoroquinolones.

In 7% of cases, the isolates without known antibiotic-resistance genes appeared to be non-susceptible (Table S4), suggesting the presence of resistance mechanisms, like loss of porins and increased efflux [22], which are not included in the ResFinder database. For example, two isolates showed intermediate susceptibility to streptomycin, and 15 isolates were non-susceptible to ampicillin/sulbactam in the absence of the genes which would explain the corresponding resistance phenotypes in those isolates. Among the tetracycline-resistant isolates, 3.3% (n=2) of isolates did not possess any known tetracycline resistance genes, while 86.9% (n=53) had a truncated version of *tet(A*) gene (1172 bp vs. 1200 bp for the full-length reference sequence from the ResFinder database). Nonetheless, the isolates without tetracycline-resistance genes or with incomplete *tet(A*) gene demonstrated a high level resistance to tetracycline (MIC >8μg/mL). This suggests the presence of additional mechanisms of resistance, undetectable by the ResFinder database.

Acquired ABR genes were frequently associated with various mobile elements. Penicillin resistance encoded by *bla*_TEM-1_ gene was associated with Tn*3* transposon located on conjugative plasmids of IncB/O/K/Z incompatibility group found in both recent SF and historical isolates (Fig. S9A) or on the putative IncI1 plasmid in several modern SJ isolates (Fig. S9B). Partial plasmid sequences were identified in historical isolates as IncB/O/K/Z and contained plasmid backbone genes encoding following products: transcriptional activator RfaH, plasmid stabilization system protein, ribbon-helix-helix protein CopG, replication initiation protein RepA, and plasmid conjugative transfer protein. This partial IncB/O/K/Z plasmid sequences in historical isolates matched analogous area in IncB/O/K/Z plasmids of recent isolates, suggesting their relatedness to each other. A determinant Tn*3*::*bla*_TEM-1_ was integrated into different loci of the IncB/O/K/Z plasmids in historical isolates in comparison with integration locus in IncB/O/K/Z plasmids of modern San Francisco cluster isolates, which suggests that dissemination of the *bla*_TEM-1_ gene is associated with the transfer via Tn*3* (Fig. S9A).

Aminoglycoside and trimethoprim resistance genes were often linked with Tn*7* transposon (Fig. S9C). A genetic region, containing an association of Tn*7* with *aadA*1 and *dfrA*1 ABR genes, as well as TDP-fucosamine acetyltransferase, and a partial sequence of Tyrosine recombinase XerC/D, was identical in recent SJ isolates and two historical samples C99 and C92, isolated in Shasta County, 2000 and Imperial County, 2002, correspondingly. However, the area upstream of Tn*7* in novel isolates showed evidence of later integration of additional Tn*7* element, leading to the loss or modification of the area containing tetracycline resistance genes found in historical isolates.

One of the historical isolates C80 (Orange County, 1987) was resistant to azithromycin and possessed macrolide resistance gene *mphA*, found in association with Tn*7* transposon on a contig which also encoded trimethoprim, sulfamethoxazole, aminoglycoside resistance, multidrug transporter EmrE, and a Mercuric resistance operon (Figure 6). Resistance to azithromycin is emerging in the United States [23], and the presence of azithromycin resistance gene in one of the isolates dating back to 1987 is remarkable.

**Figure 6.**
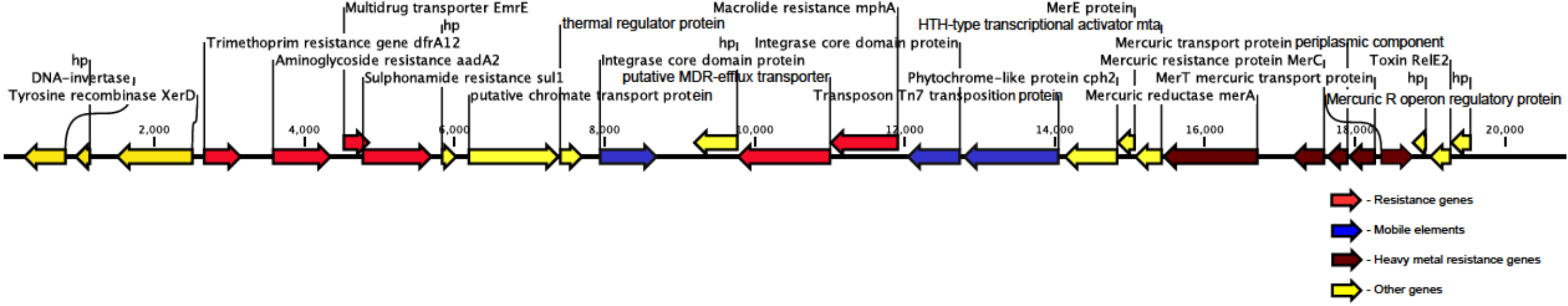
Organization of genetic surrounding of macrolide resistance gene in historical *S. sonnei* isolate C80 (Orange County, 1987) Abbreviation: hp-hypothetical protein.

### Fluoroquinolone resistance

All modern isolates from the SD/SJ population and historical *S. sonnei* isolates were susceptible to fluoroquinolones (FQ), however few historical isolates possessed several types of point mutations leading to AA substitutions in the genes encoding FQ targets-the A subunit of DNA-gyrase (GyrA) and the C subunit of topoisomerase IV (ParC) (Figure 5, Table S4) The quinolone resistance-determining region (QRDR) [24] of the GyrA in two CA historical isolates had a single amino acid substitution S83L. *S. sonnei* isolates with single mutation S83L have been previously reported to exhibit high minimum inhibitory concentration (MIC) of quinolones (nalidixic acid), but low MIC of fluoroquinolones (MIC range of ciprofloxacin 0.125-0.25μg/ml) [25]. In agreement with previous data, historical CA *S. sonnei* isolates with single mutation S83L remained phenotypically susceptible to ciprofloxacin with wild-type MIC values of ≤0.5 μg/mL. Two FQ-susceptible historical isolates had single point mutation V533A in the ParC, and one FQ-susceptible isolate had combination of ParC-V533A and GyrA-D483E mutations; none of the mutations were located in QRDR regions of corresponding proteins and were not associated with a resistant phenotype.

In contrast, all SF population isolates implicated in the outbreak expressed high level FQ-resistance. All isolates from the contemporary San Francisco outbreak had a combination of point mutations S83L and D87G in QRDR GyrA and substitution S80I in QRDR ParC, which caused a significant increase in ciprofloxacin MIC to ≥4 μg/mL (Table S4). The modern nonoutbreak isolate C42, belonging to the SF population, was phenotypically susceptible to FQ and possessed a single QRDR GyrA mutation S83L, which represents a prerequisite mutation allowing for one-step development of high-level FQ resistance if combined with another QRDR mutation [26], thus suggesting a concurrent accumulation of FQ resistance mutations in the SF population of *S. sonnei* (Figure 5).

## DISCUSSION

From our point of view, the key results of our study were: 1) the first large scale whole genome sequencing and analysis of *S. sonnei* strains from North America, 2) the delineation of two distinct yet related populations of *S. sonnei* that caused large outbreaks of shigellosis in California [SD and SJ population and SF population], 3) linkage of SD/SJ and SF populations to an older lineage of *S. sonnei* documented in California as early as 1986. This historical CA lineage was characterized as sequence type 152 and belonged to a global Lineage III with origin in Europe, 4) evidence of stx1-encoding bacteriophage in the SD/SJ population. The phage was related to the phage from pandemic *E. coli* O104:H4 with the STX-operon identical to those in *S. flexneri* and *S. dysenteriae*, 5) evidence of fluoroquinolone resistance in SF population via accumulation of mutations in the genes of antibiotic targets-GyrA and ParC.

According to the whole-genome hqSNP phylogeny, the isolates from outbreaks of 2014 - 2015 were divided into two distinct SD/SJ and SF populations. The isolates from SD and SJ epidemiological groups were assigned to two separate clusters according to PFGE profiling [9], however based on WGS we showed that SD and SJ clusters were a part of the same SD/SJ outbreak population. Many recent publications have highlighted superior discriminatory power and predictive value of WGS over PFGE for genotyping of bacterial strains [27].

Comparison to global *S. sonnei* clones placed both modern SD/SJ and SF populations as well as majority of historical CA *S. sonnei* isolates dating back as far as 1986 into the Lineage III defined by Holt et al. [13] (Figure 1A, Fig. S1). All STX-positive SD/SJ isolates clustered with Global III clade defined by Holt et al. within Lineage III, which was shown to be particularly successful at global expansion [13]. Remarkably, the isolates from SD/SJ population, as well as from SF population, were closest to the isolate from the UK, 2008, while they did not cluster with geographically proximate isolate from Mexico, which was only related to few historical CA isolates. Accepting the hypothesis that all original *S. sonnei* lineages diversified from Europe and then were introduced to other countries where they underwent localized clonal expansions [13], our findings suggest that while strain exchange with Mexico had happened in the past, the lineage ancestral to the current SD/SJ and SF *S. sonnei* populations was introduced at some point from Europe, and diversified within CA. The data suggests that the acquisition of stx1 gene happened after the original Lineage III *S. sonnei* clone had already spread in CA. The contemporary C130 and historical C84 STX-negative *S. sonnei* isolates, which both clustered with STX-positive SD/SJ outbreak isolates, most likely represent an ancestral STX-negative lineage for the SD/SJ population, prior to STX-bacteriophage acquisition. It appears that global *S. sonnei* clones belonging to Lineages I, II, and III were introduced to CA independently during the last three decades.

All recent *S. sonnei* outbreak isolates as well as all but one historical isolates belonged to a sequence type ST152. This long-term persistence of a single clone in CA over three decades is remarkable, and could indicate that ST152 is a very successful *S. sonnei* clone. There were previous examples when the same sequence types persisted in the same geographical area for a long time, e.g. ST245 in China was found in 1983 as well as 26 years later in 2009 [12]. This phenomenon, however, is most likely explained by a low resolution of MLST for subtyping of clonal *S. sonnei* population, which has been noted before [28].

Isolates from the SD/SJ *S. sonnei* population, which caused outbreak of severe diarrheal disease [9], carried stx1 genes encoded by a novel lambdoid bacteriophage in all of the isolates. The STX-bacteriophage from *E. coli* O104:H4 strain implicated in an European outbreak of 2011 was found to be the closest known relative to STX-phage of SD/SJ *S. sonnei* isolates based on overall similarity of the whole-phage sequences. This outbreak was one of the world’s largest outbreaks of food-borne disease in humans with 855 HUS and 53 fatalities [29, 30]. Considering the particular proximity of CA STX-phage to the phage from *E. coli* O104:H4 strain 2011C-3493 introduced from Germany to the US in 2011, we speculate that introduction of STX-phage into the SD/SJ *S. sonnei* population happened after the outbreak *E. coli* O104:H4 was brought to the US, followed by a local diversification of STX-phage, which involved recombination of STX-operon to yield the final phage with general architecture of *E. coli* phage and STX-operon from *S. flexneri* or *S. dysenteriae*. However, it is also possible that STX-phage from pandemic *E. coli* O104:H4 underwent the modifications via recombination with other European phages, prior to its introduction to CA *S. sonnei* clone. The transfer of STX-bacteriophage is the only gene exchange event we detected between *E. coli* O104:H4 and *S. sonnei* genomes. It would be pertinent to emphasize that the German outbreak strains of *E. coli* also contained distinct STX operon with *stx2* gene, several virulence and antibiotic-resistance elements [31].

It has been demonstrated previously that phage phylogeny should be inferred from a combination of protein repertoires and phage architecture rather than from a single gene (e.g. integrase) sequence [32, 33]. The mosaic structure of the lambdoid phages poses a limitation for such single locus-phylogeny approach. Thus, even though the integrase phylogeny showed that CA isolate has an integrase gene more closely related to older Georgian than with recent German *E. coli* O104:H4 isolate, this does not contradict the relatedness of CA phage to the outbreak lamboid-phage shown based on the cumulative data derived from the whole phage sequence. This also could be explained by a long-term persistence of phages belonging to a given Integrase family in Europe.

STX-1 encoding bacteriophage was not found in the isolate C130 even though it was closely related to SD/SJ *S. sonnei* population. It is possible that the bacterial population as a whole acquired STX1-encoding bacteriophage more recently, and concurrently. Such a scenario would also suggest that the increase in the virulence took place either by the spread of STX-positive clones or by the horizontal transfer of stx1- genes by bacteriophages. The potential of bacteriophages to transfer STX genes from one *S. sonnei* strain to another has been described previously [3]. Not all the *S. sonnei* strains have a suitable genetic background to be able to express STX efficiently [34]. The modern STX-positive SD/SJ population of *S. sonnei* was shown to be able to express STX and thus is proven to possess the genetic background which is adapted for the increased virulence. The adaptation of SD/SJ *S. sonnei* genomic background to retain and express STX stably by itself is a prerequisite for future emergence of more virulent strains in CA. Another example of SD/SJ population increasing its virulence is an acquisition of <u>i</u>ncreased <u>s</u>erum <u>s</u>urvival virulence gene iss, which occurred seemingly recently in evolution of SD/SJ S. sonnei clone. To our knowledge, this is the first report of increased serum survival determinant (iss) being found in *Shigella* species.

Even though the treatment of infection caused by *S. sonnei* with antibiotics is not a standard procedure, it is indicated for the treatment of severe cases in order to reduce duration of symptoms [35]. Antibiotic resistance of *S. sonnei* is however on the rise [10, 36, 37]. This highlights the importance of ABR monitoring in *S. sonnei* populations. We detected resistance markers to multiple classes of antimicrobials in CA *S. sonnei*. 91.2% of modern isolates and 81.8% of historical isolates were multi-drug resistant (MDR) according to the definition of MDR microorganisms as demonstrating “non-susceptibility to at least one agent in three or more antimicrobial categories” [38]. Acquired ABR genes in CA *S. sonnei* were frequently associated with various mobile elements and conjugative plasmids, likely contributing to their dissemination. For example, MDR transposon Tn*7* has been shown to be a crucial part of ABR evolution in the local populations of *S. sonnei* Lineage III [13]. Once acquired, ABR genes tend to stay in the bacterial population, e.g. traditional antibiotics like cotrimoxazole and tetracycline are no longer in use for patient treatment [12], but the ABR genes still persist in the modern SD/SJ and SF populations of the CA *S. sonnei*. Particularly worrisome was the fluoroquinolone resistance detected in all of the isolates belonging to a modern SF *S.sonnei* population, implicated in the outbreak. FQ-resistance in all SF outbreak isolates was mediated by a combination of double mutation in QRDR GyrA of DNA gyrase and one mutation in QRDR ParC of topoisomerase IV. All historical isolates as well all other modern isolates were susceptible to FQ, however few of those phenotypically susceptible CA *S.sonnei* isolates possessed a prerequisite single AA substitution in QRDR of the GyrA, including two isolate belonging to CA Lineage III: one historical isolate from 2007 and one modern non-outbreak isolate belonging to SF population. Single mutations in QRDR GyrA were shown previously to confer a low level of FQ resistance and to serve a prerequisite for further resistance escalation via stepwise mutations acquisition [26]. Therefore, the data suggests that evolution of SF population from ancestral *S. sonnei* lineage is occurring via increase in fluoroquinolone resistance mediated by the accumulation of FQ-resistance mutations.

There are certain limitations of this study: 1) temporal and spatial representation was sporadic in *S. sonnei* strains in our culture collection, 2) the short read sequencing by synthesis used to generate data might limit complete assembly of *S. sonnei* genomes and enumeration of genome-wide SNPs, 3) no plasmid transformation followed by re-sequencing could be performed to confirm postulated genetic exchanges among *S. sonnei* populations. Although the direct experimental evidence is lacking, Shiga-toxin production and other virulence elements discovered in SD/SJ population appeared to be among the contributors that lead to serious manifestations of gastrointestinal disease in CA shigellosis outbreak including bloody diarrhea in 71% of patients [9]. Fortunately, there were no fatalities and none of the affected patients developed more serious hemolytic uremic syndrome (HUS). One potential clue to the absence of severe manifestations could be that Shiga-toxin positive *S. sonnei* strains contained Stx1. The toxins encoded by Stx1 and Stx2 are known to elicit variable pathology among affected individuals. A number of investigators have demonstrated that *E. coli* O157 Shiga-toxin producing strains carrying Stx2 caused more severe disease including HUS than Stx1 positive strains [39–41].

Conclusion: Two distinct populations of *S. sonnei* (SD/SJ and SF) have been delineated in the recent shigellosis outbreaks in California. These populations evolved from a common lineage of *S. sonnei* likely present in California as early as 1986. The suggested evolutionary pathways were: 1) increased virulence via acquisition of a phage from the *E. coli* O104:H4 German outbreak strain and STX-operon from *S. flexneri* or *S. dysenteriae*, and 2) emergence of fluoroquinolone resistance via the accumulations of point mutations in *gyrA* and *parC* genes. The modern CA *S. sonnei* populations were related to the global Lineage III, which originated in Europe and was known for its successful expansion around the world. Thus, the CA *S. sonnei* lineage continues to evolve by the acquisition of increased virulence and antibiotic resistance, and enhanced surveillance is advocated for its early detection in future shigellosis outbreaks.

## MATERIALS AND METHODS

### Isolates

Sixty-eight *S. sonnei* human isolates from CA (57 outbreak-related from 2014-2015 and 11 archival isolates from 1980-2008), were identified and serotyped by standard methods [42]. PCR-detection of *stx*_*1*_ and *stx*_*2*_ genes, and Vero cell neutralization assay for confirmation of STX-production were performed as previously described [43]. A list of isolates is presented in the Table S1. A map with geographical distribution of the location of origin of the isolates is in Figure 1B.

### WGS and data analysis

DNA was extracted with a Wizard Genomic DNA Kits (Promega, Madison, WI). Sequencing libraries were constructed using the Nextera XT (Illumina Inc., San Diego, CA) library preparation kits. Sequencing was performed using 2 × 300 bp sequencing chemistry on an Illumina MiSeq Sequencer as per manufacturer's instructions.

For genome-wide SNP identification, paired-end reads were mapped to the reference genome of *S. sonnei* Ss046 (NC_007384.1) with masking of the mobile and phage elements and phylogenetic tree was built using CLCbio Genomic Workbench 8.0.2 (Qiagen, Aarhus, Denmark). A phylogenetic tree was generated using maximum likelihood phylogeny (under the Jukes-Cantor nucleotide substitution model; with bootstrapping) based on high-quality single nucleotide polymorphisms (hqSNPs). SNPs were called in coding and non-coding genome areas using SAMtools mpileup (v.1.2; [44]) and converted into VCF matrix using bcftools (v0.1.19; http://samtools.github.io/bcftools/). Variants were parsed using vcftools (v.0.1.12b; [45]) to include only high-quality SNPs (hqSNPs) with coverage ≥5×, minimum quality > 200, minimum genotype quality (GQ) 10 (--minDP 5; --minQ 200; --minGQ 10; --remove-indels), with InDels and the heterozygote calls excluded.

Phylosift [15] was used to (1) identify 37 “universal” genes in a set of genomes (including those generated here) (2) generate alignments for each gene family and then concatenate the alignments. A phylogenetic tree was inferred from the concatenated alignments using RAxML 7.2.6. bipartition trees were generated using 1000 bootstraps. COG-and Pfam-based identification and clustering was done using the DOE Joint Genome Institute (JGI) Integrated Microbial Genomes (IMG) system (https://img.jgi.doe.gov/cgi-bin/mer). The presence or absence matrices were fed to RAxML 7.3.0 [46] to produce a best tree from 50 boostraps using the gamma model rate of heterogeneity for binary input. COG abundance profile and gene homologs search was performed using JGI IMG tools.

*De novo* assembly for each genome was done on CLCbio GW8.0.2. The assembled genomes of *S. sonnei* isolates had 30-159 × sequencing coverage. Genomes were annotated with prokka v1.1, the JGI IMG database, the Center for Genomic Epidemiology (CGE) (ResFinder, VirulenceFinder, PlasmidFinder) [47], and the Phage Search Tool (PHAST) [48] online resources. *In silico* multi-locus sequence typing (MLST) was performed using the CGE online tool [11] against the MLST database *E. coli#*1 [49]. To estimate similarity between the bacteriophages, sequence alignment and a neighbor-joining tree (based on the estimate of the shared gene content) were generated using the progressiveMauve program [50]. Additionally, sequences of integrases (both the DNA for the genes and the encoded amino acids for the proteins), derived from the sequences of STX-phage belonging to *Shigella* spp. and *E. coli* strains, were aligned using CLCbio GW8.0.2. general aligner and a neighbor-joining tree was generated as above.

Recombination test was performed on genome-wide hqSNP alignment using Recombination Detection Program v. 4.67 (RDP4) [14]. The following algorithms included into the RDP4 package were applied to search for the recombination events: RDP, BOOTSCAN, GENECONV, MAXCHI, CHIMAERA, SISCAN, 3SEQ, PHYLPRO, and VisRD. A window size of 100 and a step size of 30 were used.

### Antimicrobial susceptibility and resistance testing

Antimicrobial susceptibility testing of *S. sonnei* isolates was performed using Microscan Dried Gram Negative panels Neg MIC 38 (Beckman Coulter, Brea, CA, USA); the minimum inhibitory concentration (MIC) results were read and interpreted according to the manufacturer's instructions. Streptomycin (10 μg) and azithromycin (15 μg) BBL Sensi-Discs (Becton Dickinson, Franklin Lakes, NJ, USA) were used to determine susceptibility to the corresponding antimicrobials. Standard quality control strains were tested in parallel as required in respective product inserts.

## SUPPLEMENTAL MATERIAL

**Figure S1.**
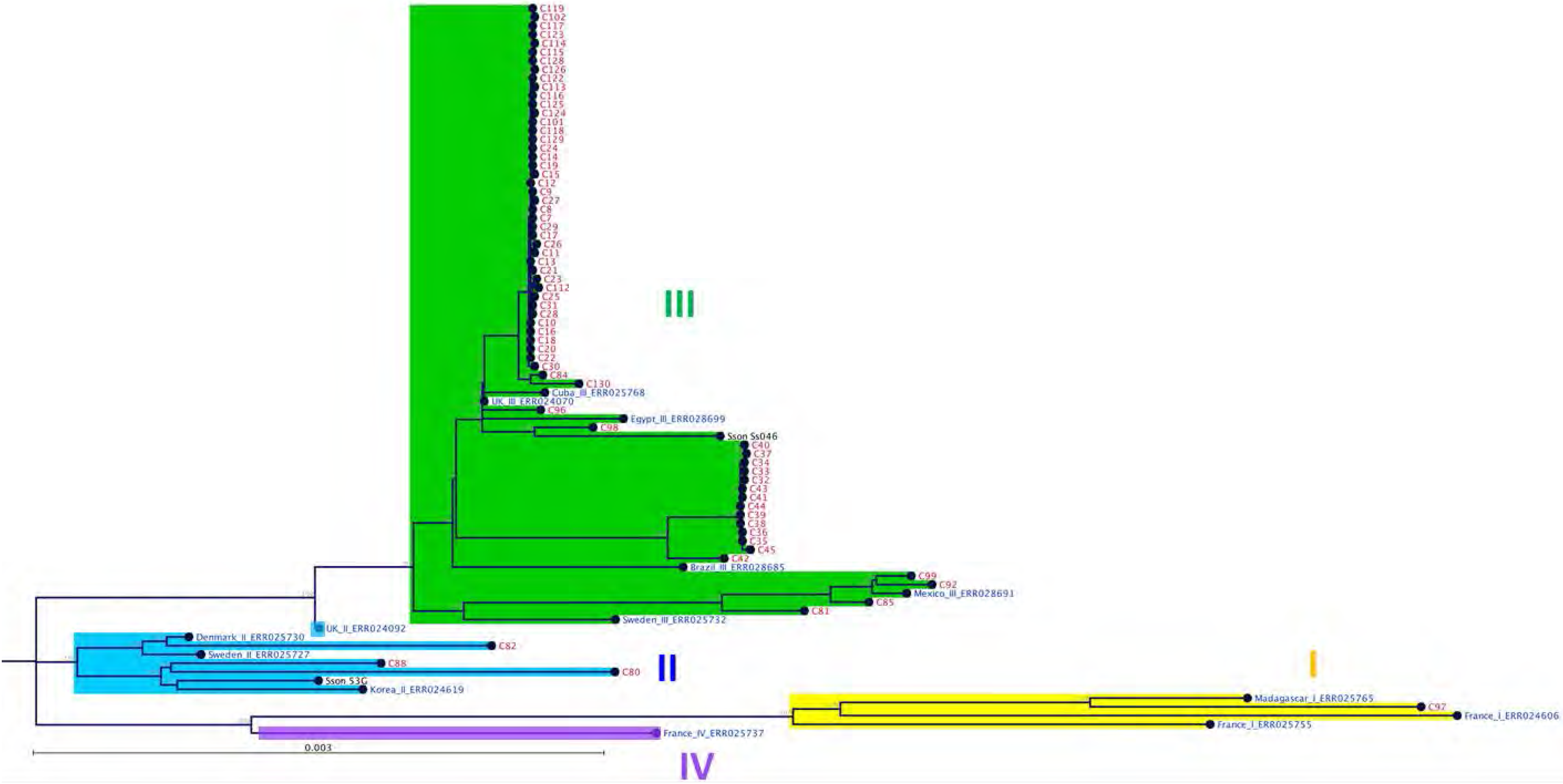
Clustering of CA *S. sonnei* isolates with *S. sonnei* strains of global lineages as per Holt et al. based on a maximum-likelihood phylogenetic tree built using genome-wide hqSNPs. Color of node labels: Red-*S.sonnei* isolates from CA; Blue-global isolates from Holt et al. publication. Branches highlight color: Yellow-Lineage I; Blue-Lineage II; Green-Lineage III; Purple-Lineage IV. Tree is rooted to *E. coli*.

**Figure S2.**
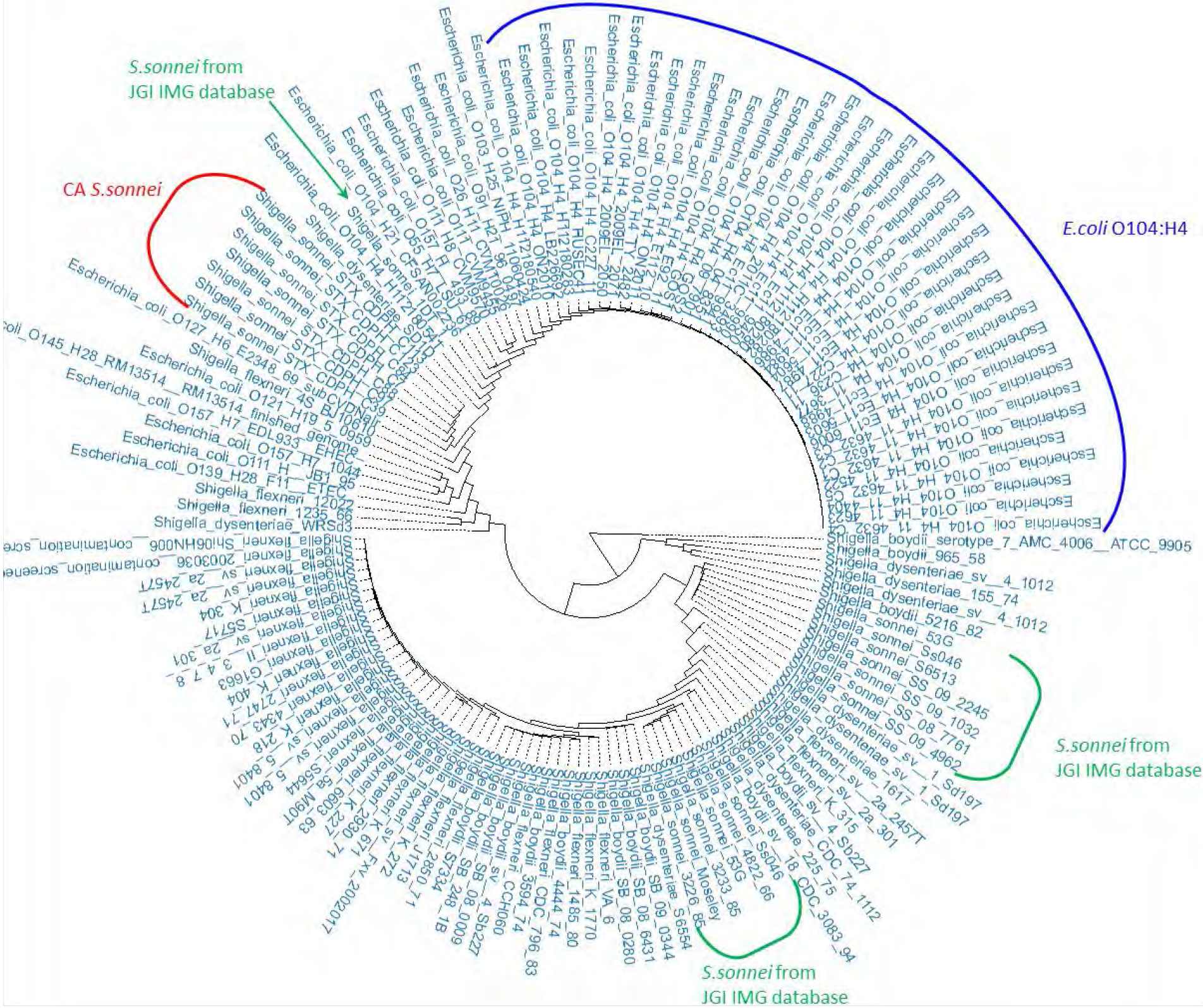
Hierarchical Clustering of CA representative *S. sonnei* isolates with *E. coli* and other *Shigella* species from JGI IMG database. A. Clustering based on COG profiles (presence/absence & abundance). Color of the brackets: Red-representative *S. sonnei* from California; Green-other *S. sonnei* from JGI IMG database; Blue-*E. coli* O104:H4 strains from JGI IMG database.

**B.**
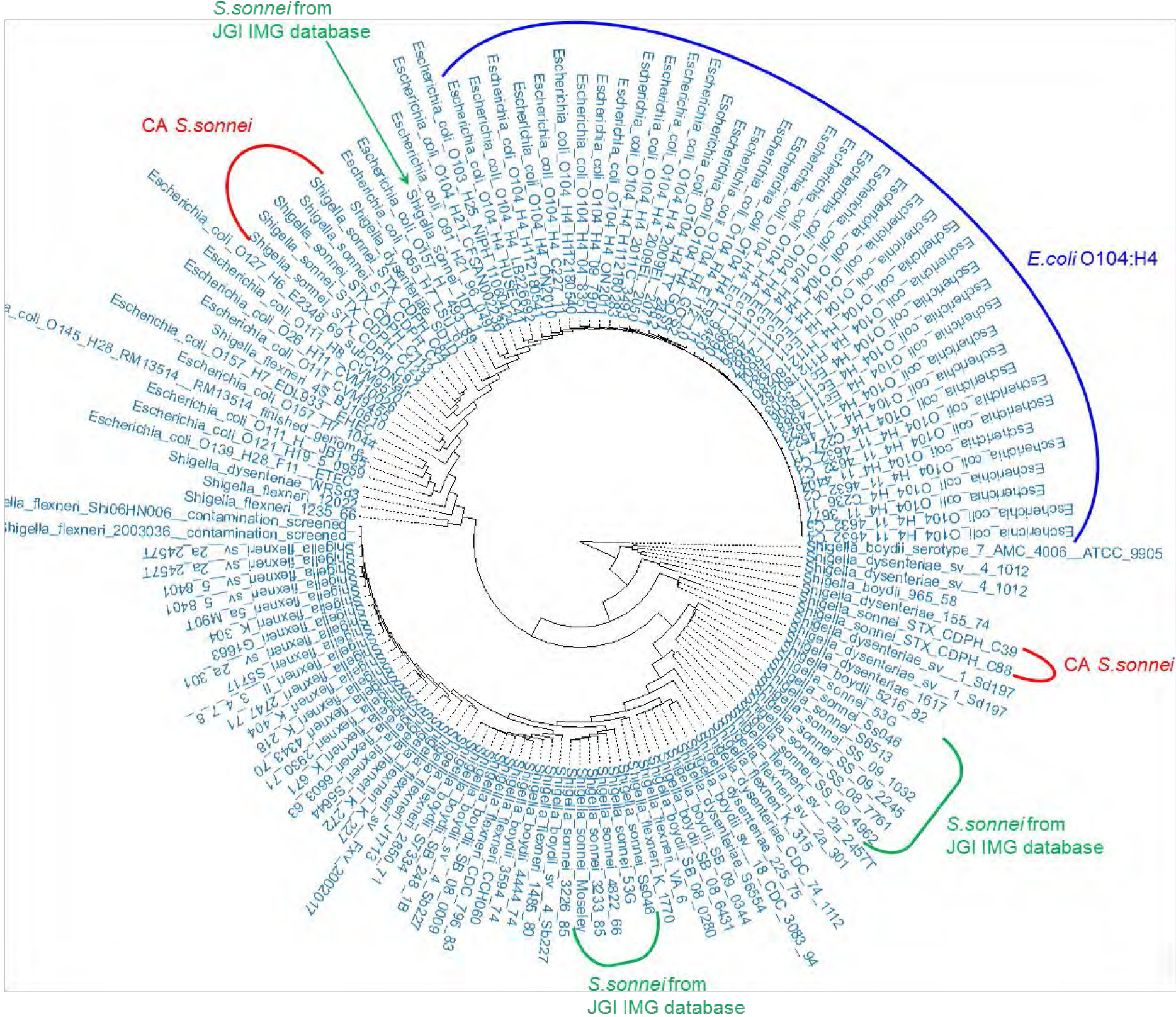
Clustering based on pfam profiles (presence/absence & abundance) Color of the brackets: Red-representative *S. sonnei* from California; Green-other *S. sonnei* from JGI IMG database; Blue-*E. coli* O104:H4 strains from JGI IMG database.

**Figure S3.**
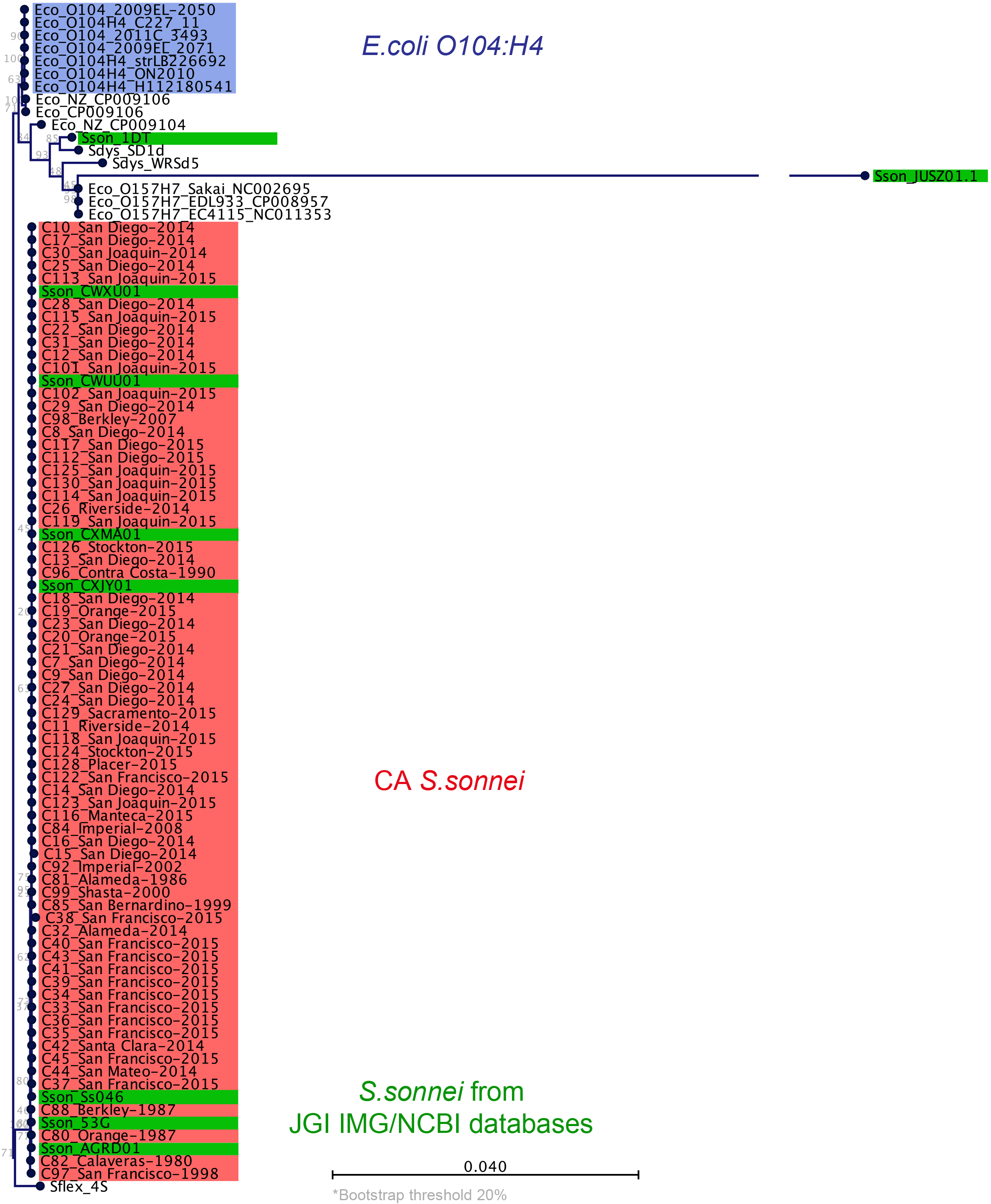
Comparison of CA *S. sonnei* with *E. coli* strains and other publicly-available genomes of *S. sonnei* based on nucleotide sequence. A. Maximum likelihood clustering from PhyloSift nucleotide-based phylogeny. Background color: Red-*S.sonnei* from California; Green-other *S.sonnei* from JGI IMG and NCBI databases; Blue-*E.coli* O104:H4 strains from JGI IMG database.Bootstrap threshold 20%.

**B.**
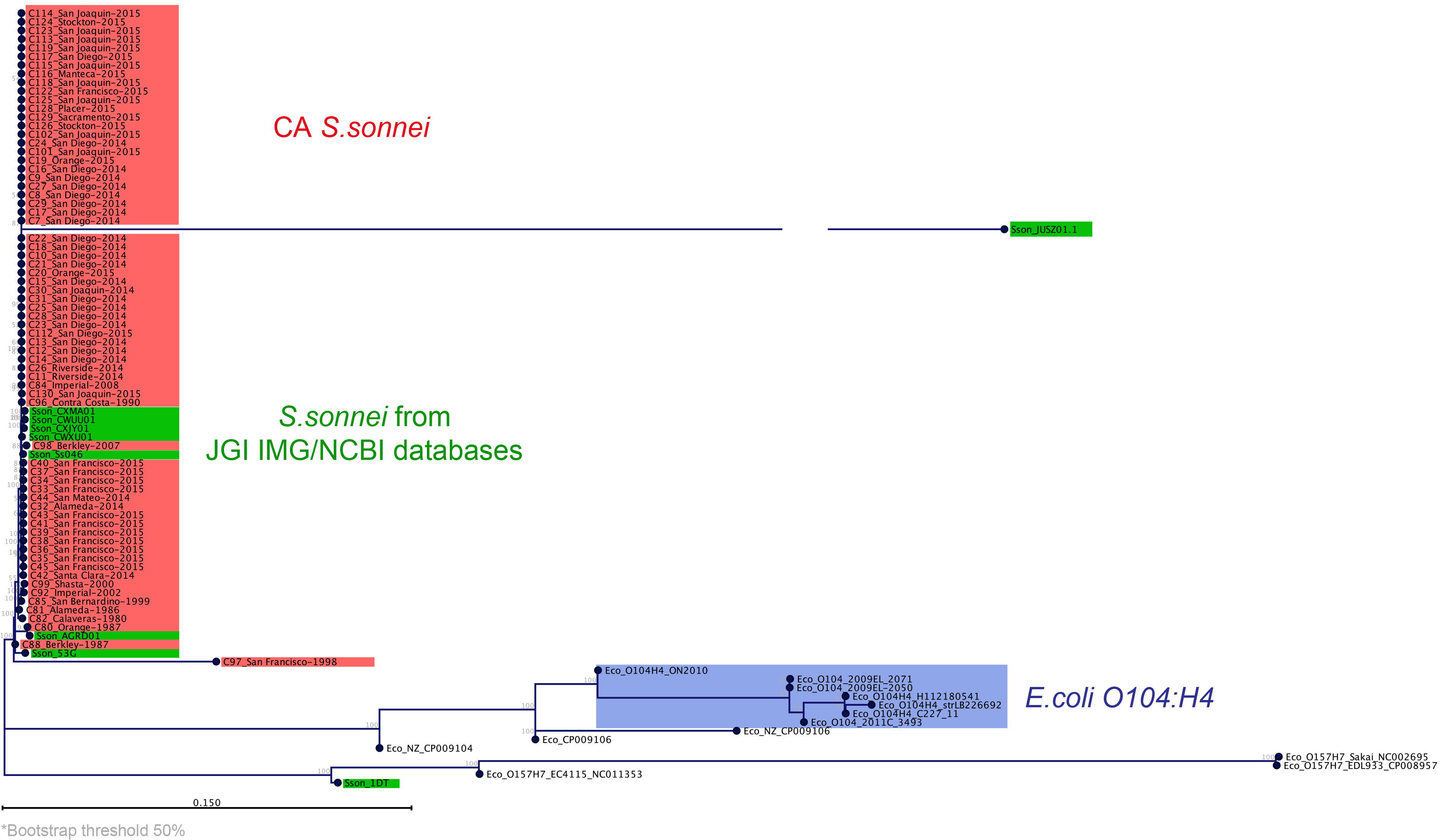
Maximum likelihood phylogeny of CA *S. sonnei, E. coli*, and publicly-available *S. sonnei* genomes based on genome-wide hqSNPs. Background color: Red-*S.sonnei* from California; Green-other *S. sonnei* from JGI IMG and NCBI databases; Blue-*E.coli* O104:H4 strains from JGI IMG database. Bootstrap threshold 50%.

**Figure S4.**
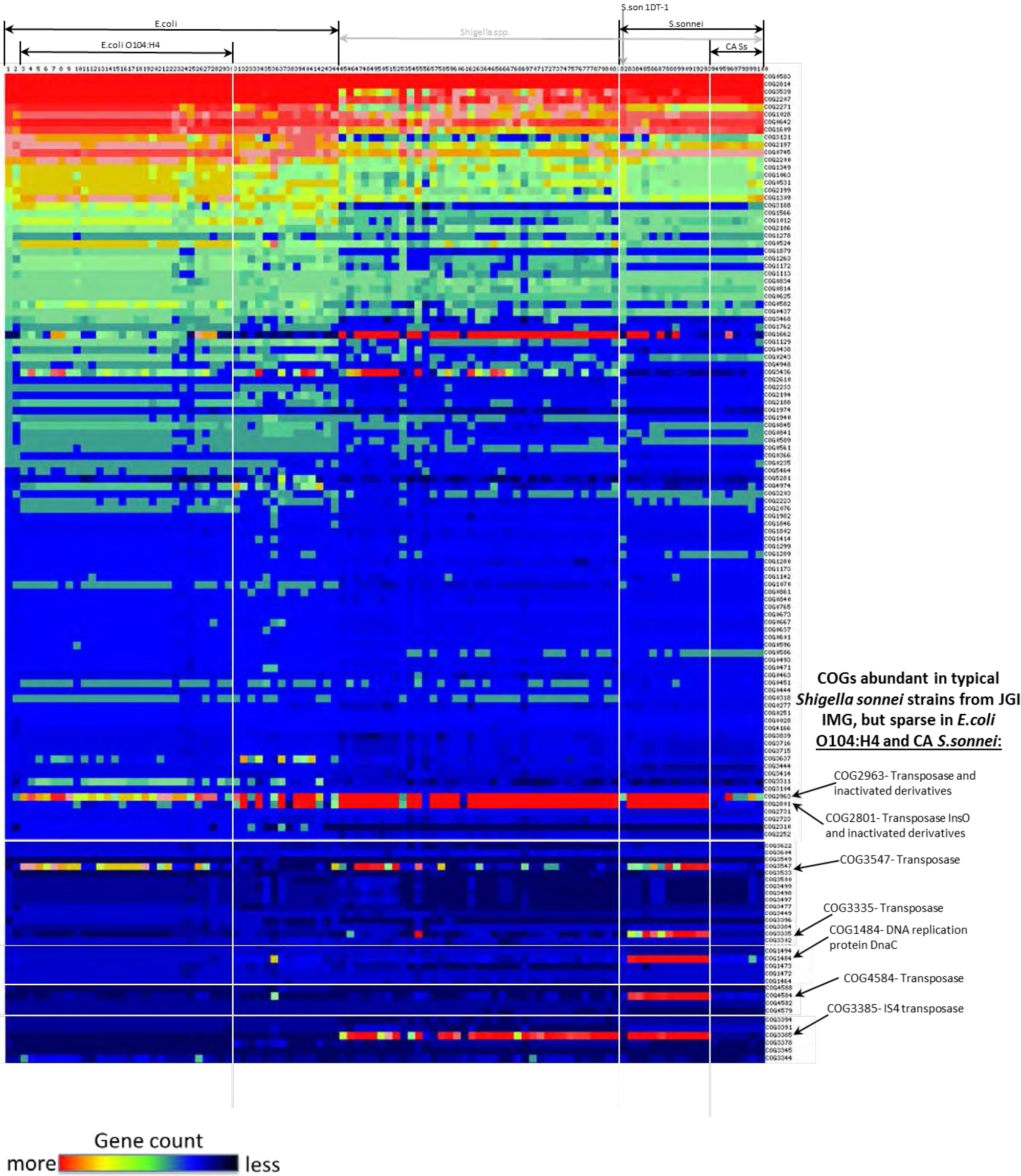
Comparison of COG abundance profiles of representative CA *S. sonnei* with *E. coli* strains and other *Shigella* species from JGI IMG database. The heat map represents gene count for different COGs.

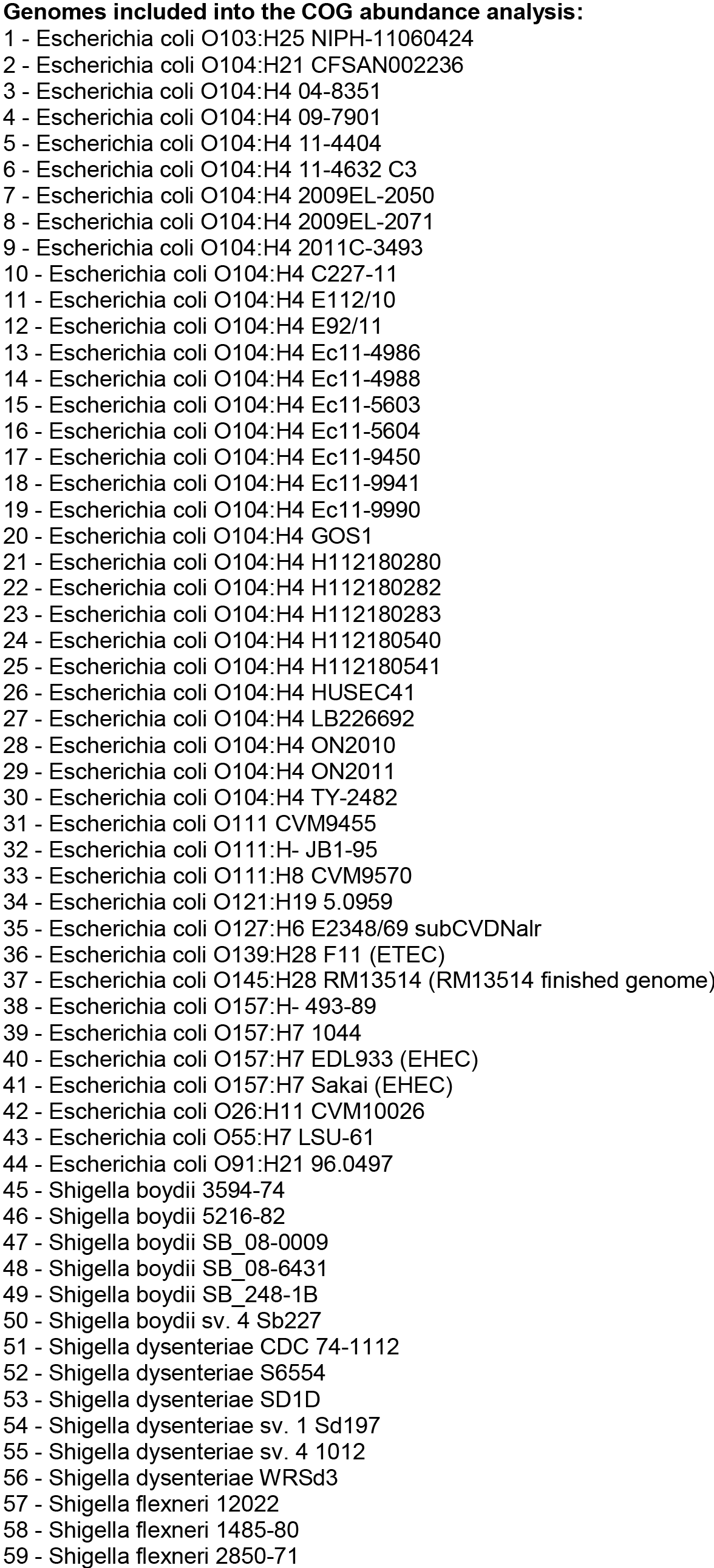

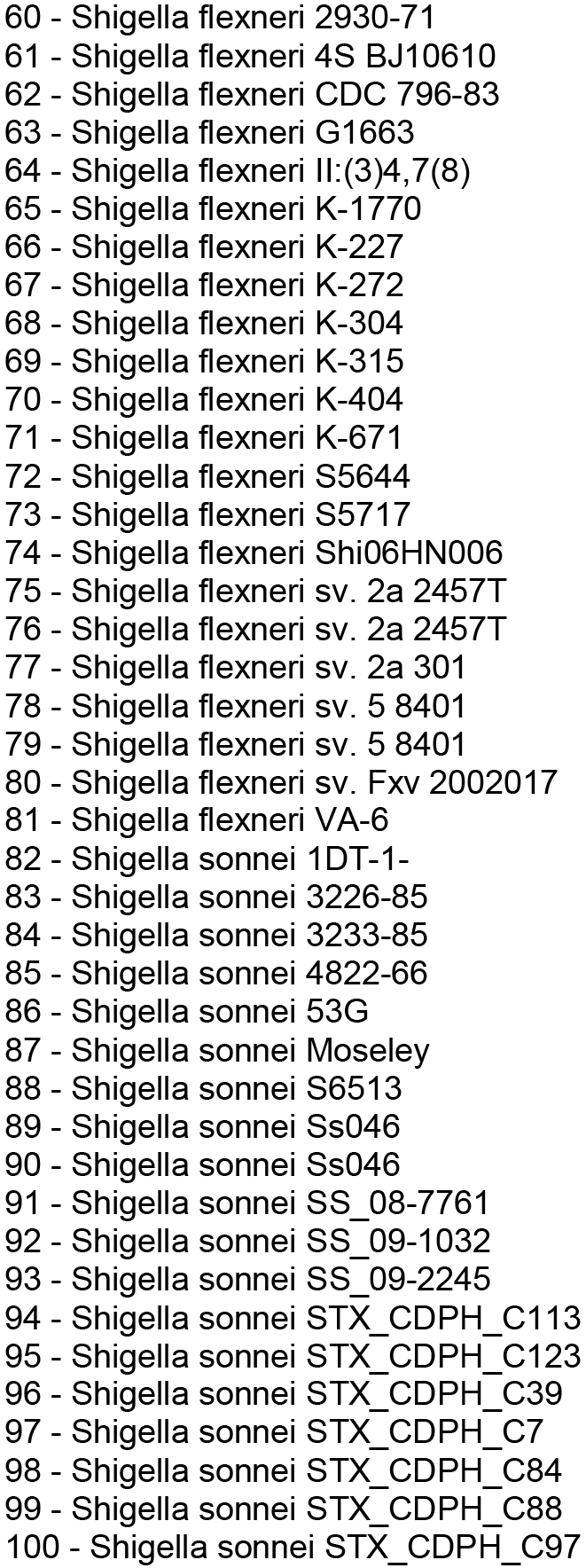

**Figure S5.**
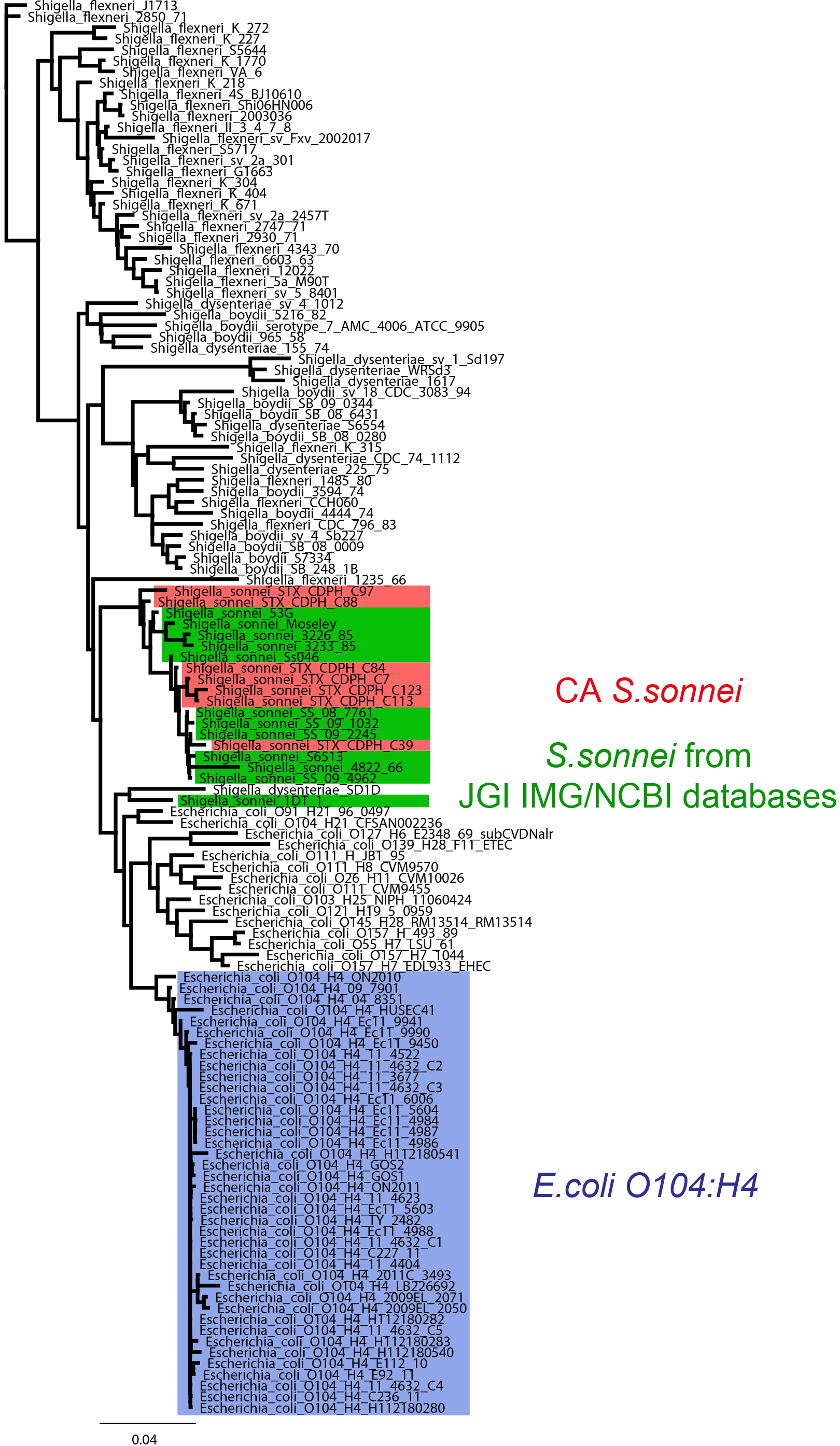
Revised pfam phylogeny. Maximum Likelihood clustering of CA representative *S.sonnei* isolates with *E.coli* and other *Shigella* species from IMG JGI database based on pfam profiles (presence/absence). Background color: Red-representative *S.sonnei* from California; Green-other *S. sonnei* from JGI IMG database; Blue-*E.coli* O104:H4 strains from JGI IMG database.

**Figure S6.**
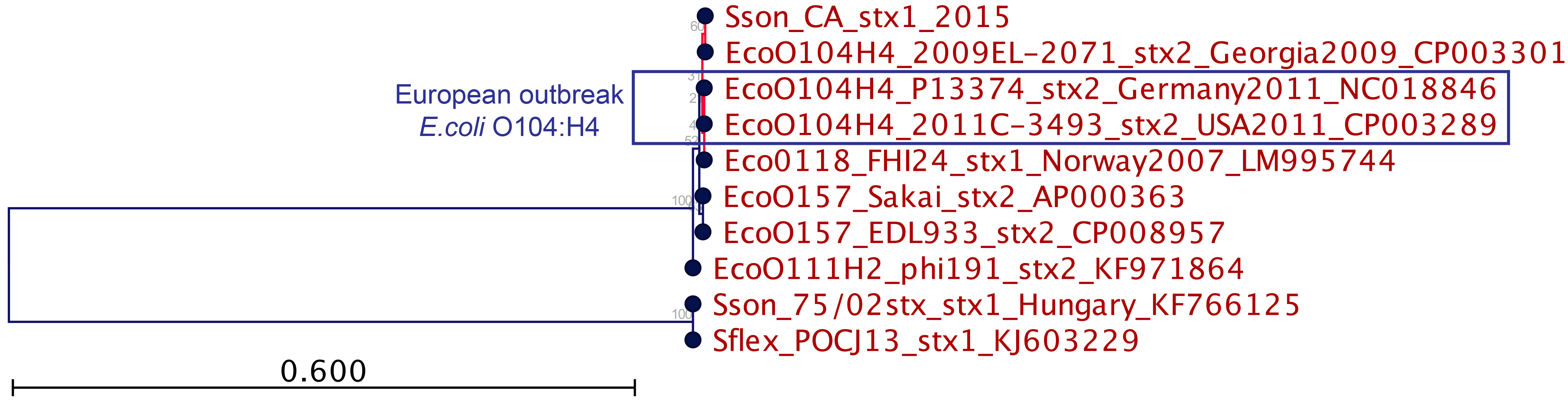
Neighbor-Joining phylogeny of the CA STX1-phage based on amino acid sequence of Integrase protein. Subtree containing CA STX1-phage is highlighted with red branch color.

**Figure S7.**
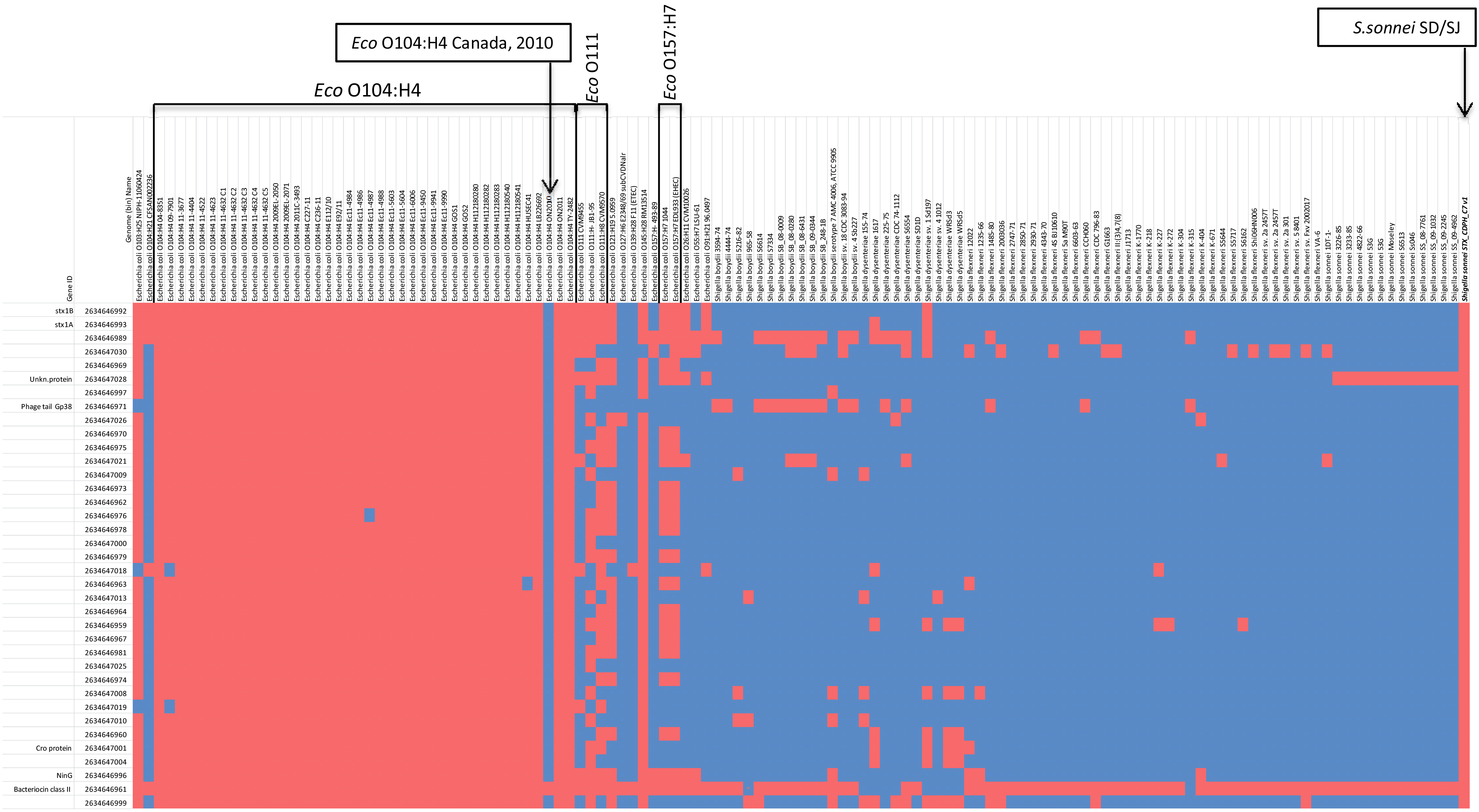
Distribution of CA STX1-bacteriophage genes in different *E. coli* and *Shigella* serovars from IMG database. Gene IDs from IMG database are listed on the left side of the graph. Color of the blocks: red - gene is present, blue - gene is absent. Min 10% ID was used as a similarity cutoff.

**Figure S8.**
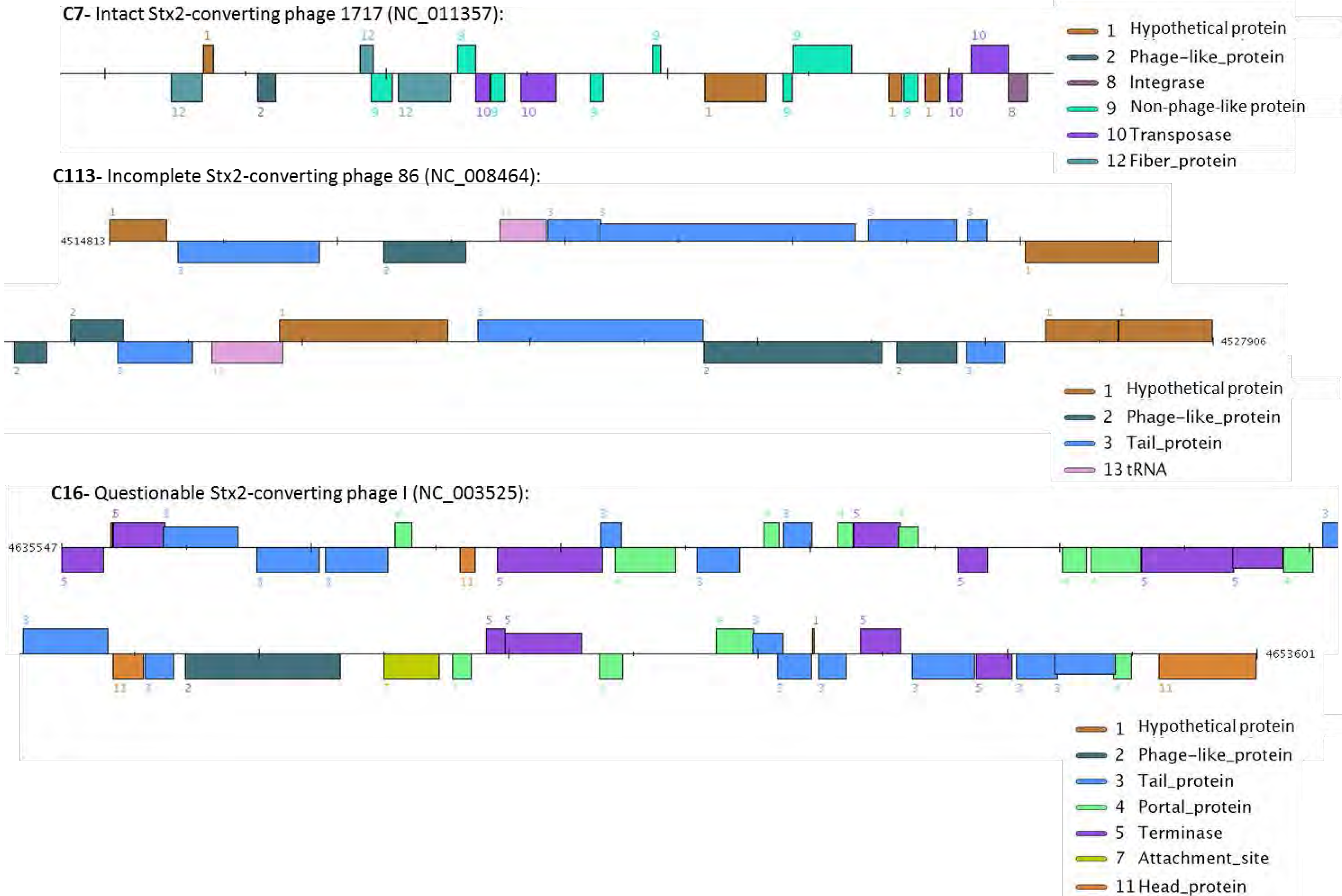
Cryptic STX-converting prophages found in CA *S. sonnei* genomes.

**Figure S9.**
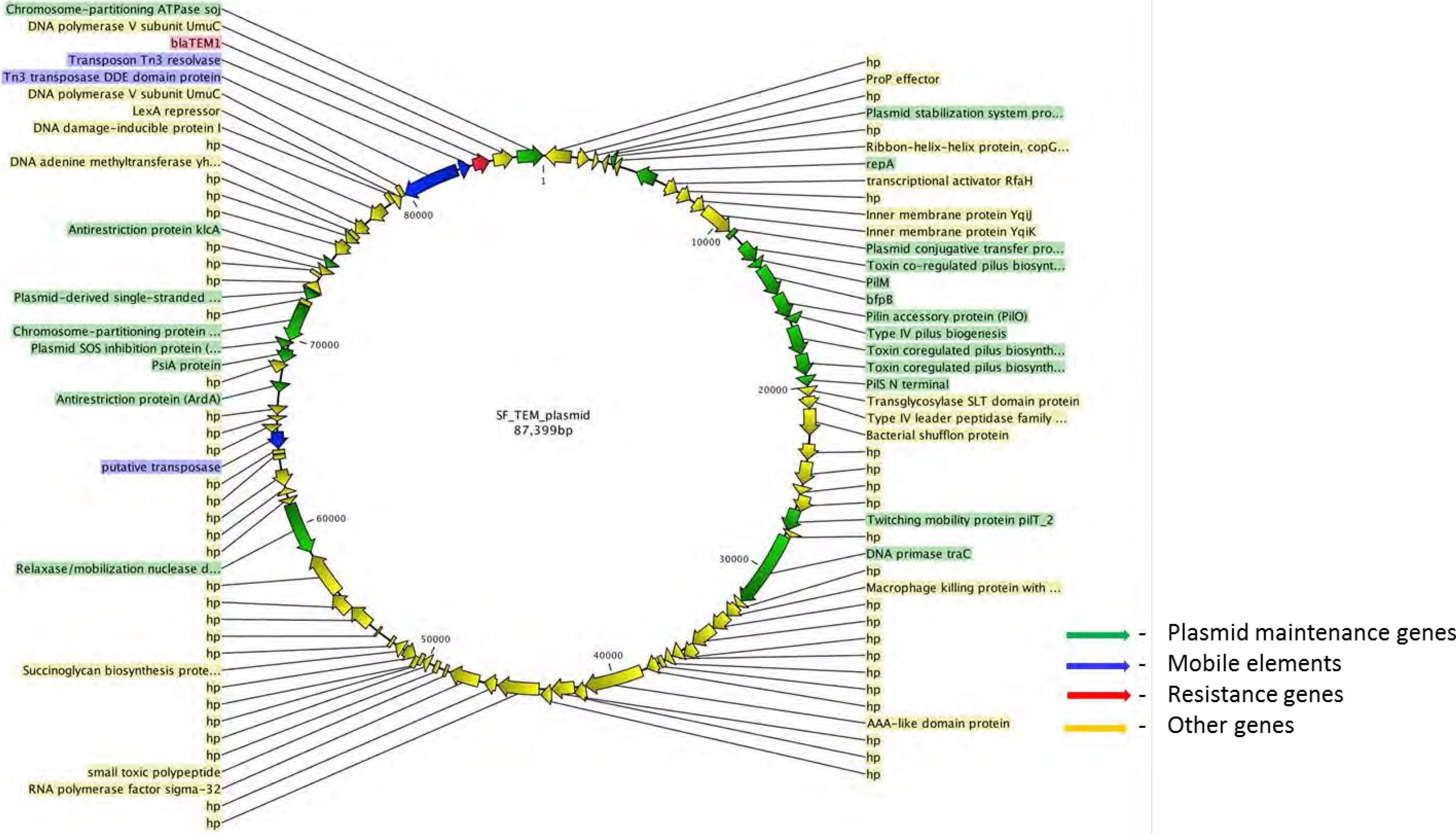
Plasmids and other mobile elements encoding antibiotic resistance in CA *S. sonnei*. A. Organization of *bla*_TEM-1_- encoding IncB/O/K/Z conjugative plasmid from modern SF population isolates

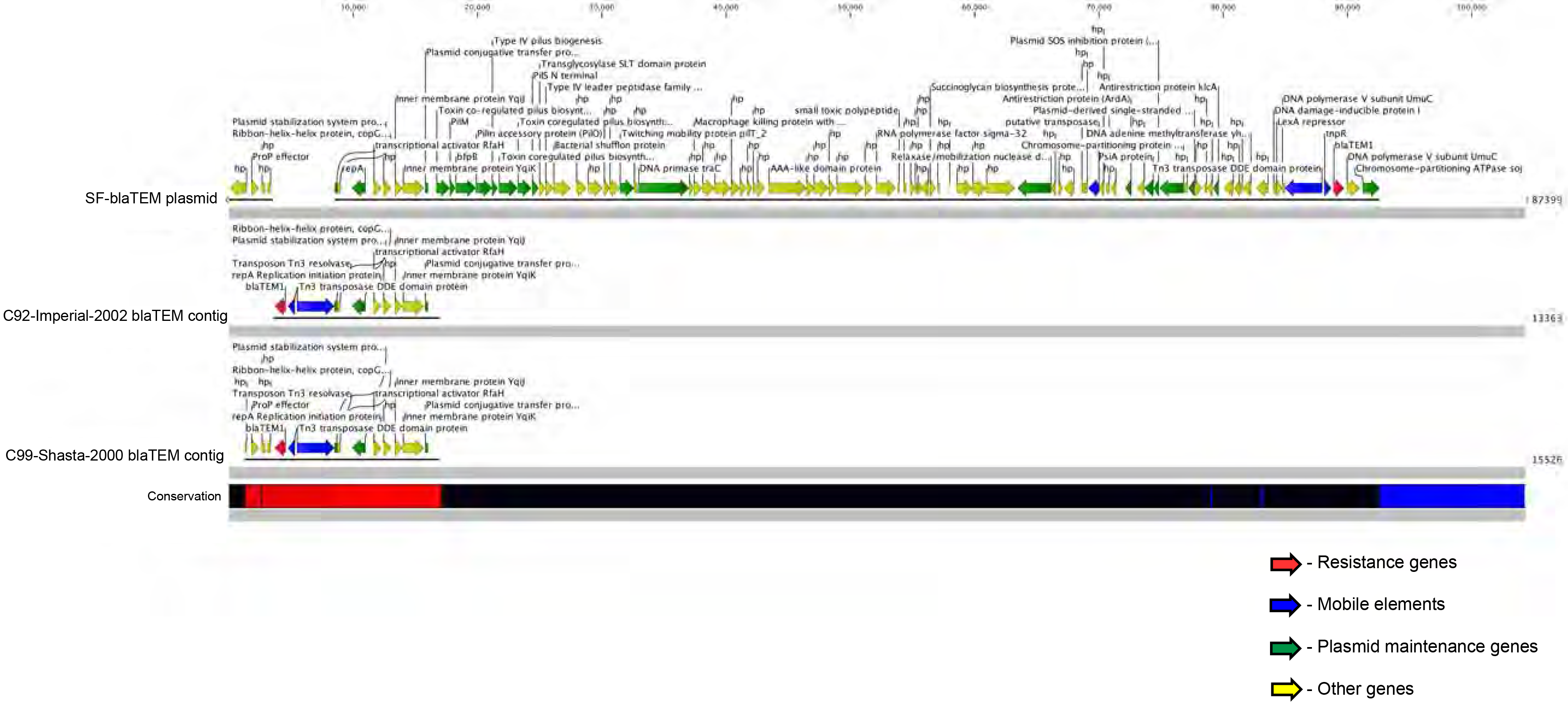
Different integration sites of Tn3::*bla*_TE_M_-1_ on IncB/O/K/Z plasmids found in recent SF isolates and in historical *S. sonnei*.

**B.**
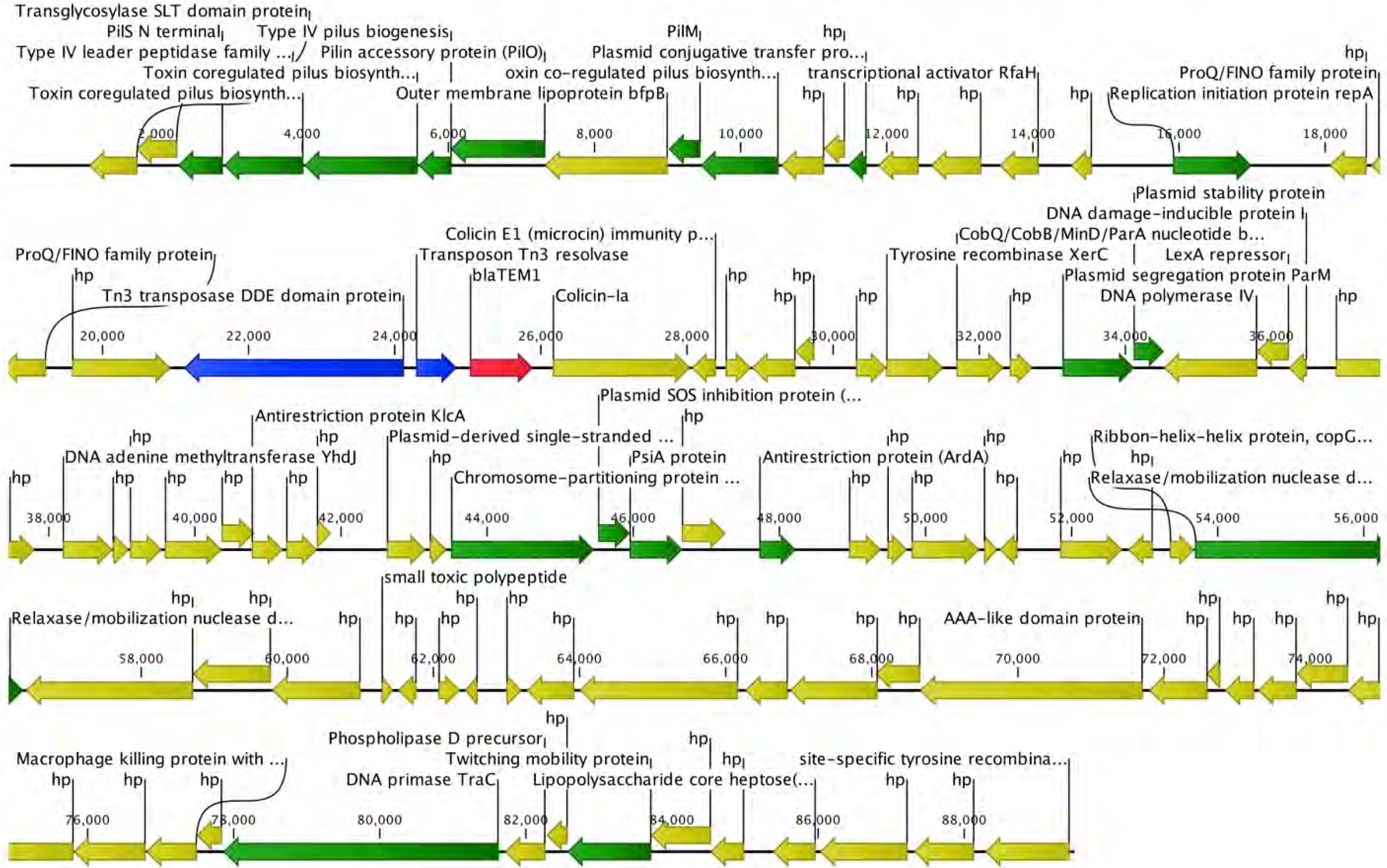
A representative genetic surrounding of *bla*_TEM-1_ gene on a putative IncI1 conjugative plasmid from a modern SJ *S. sonnei* isolate.

**C.**
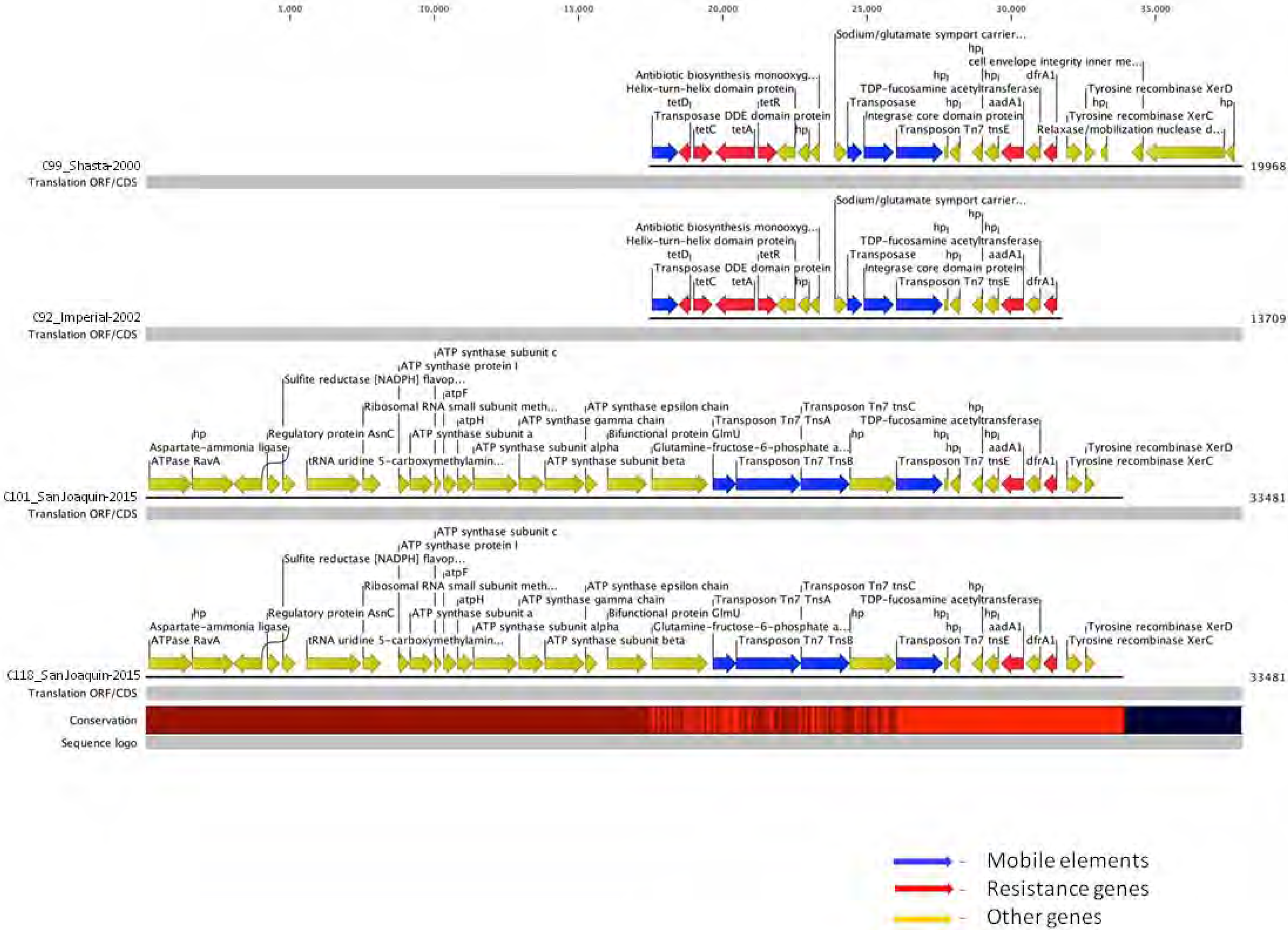
Representative organization of *sul2-strA-strB-tetA* resistance genes cluster in SD/SJ and historical CA *S. sonnei* isolates.

**Supplementary Table 1.**
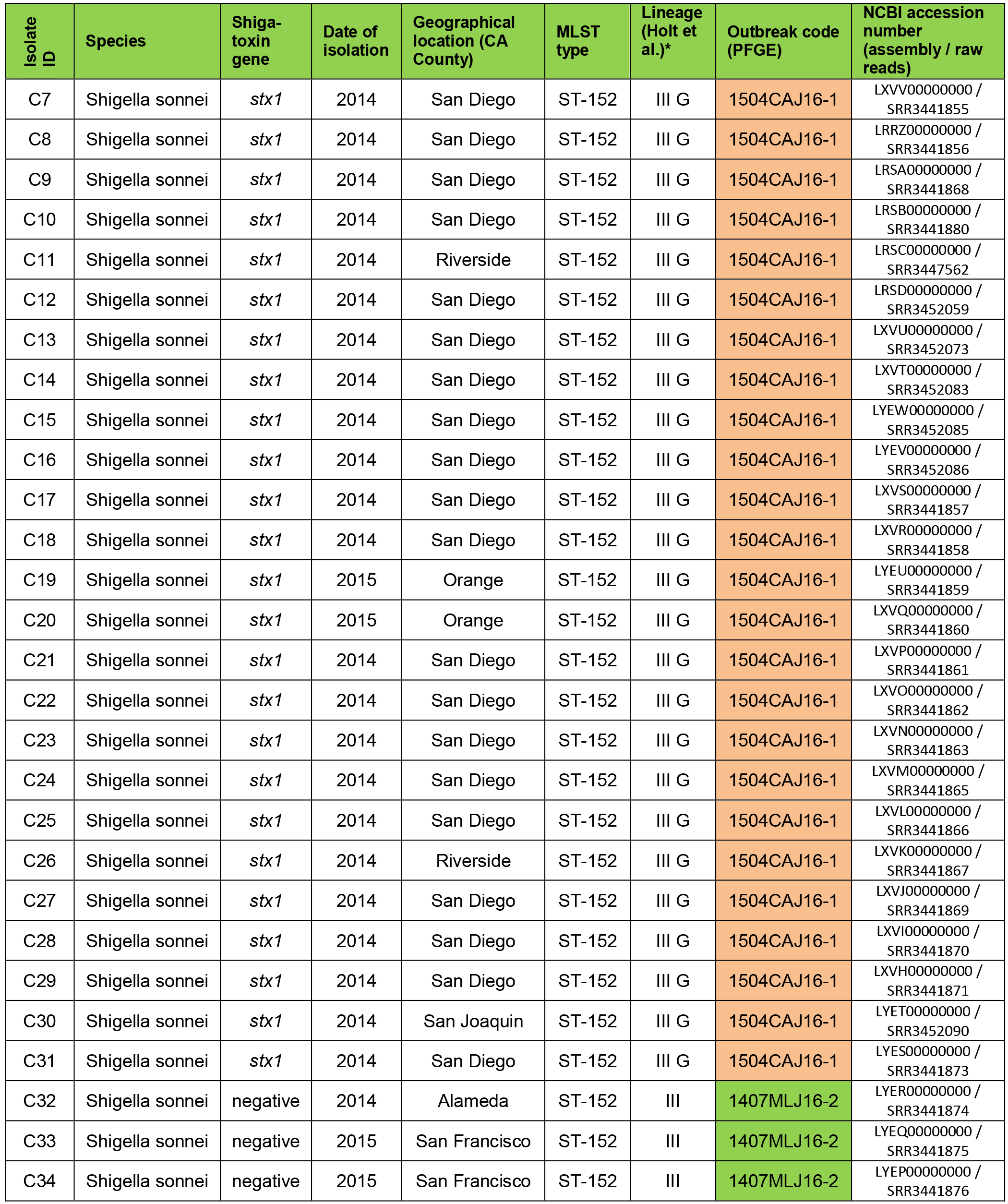
List of CA *S. sonnei* isolates

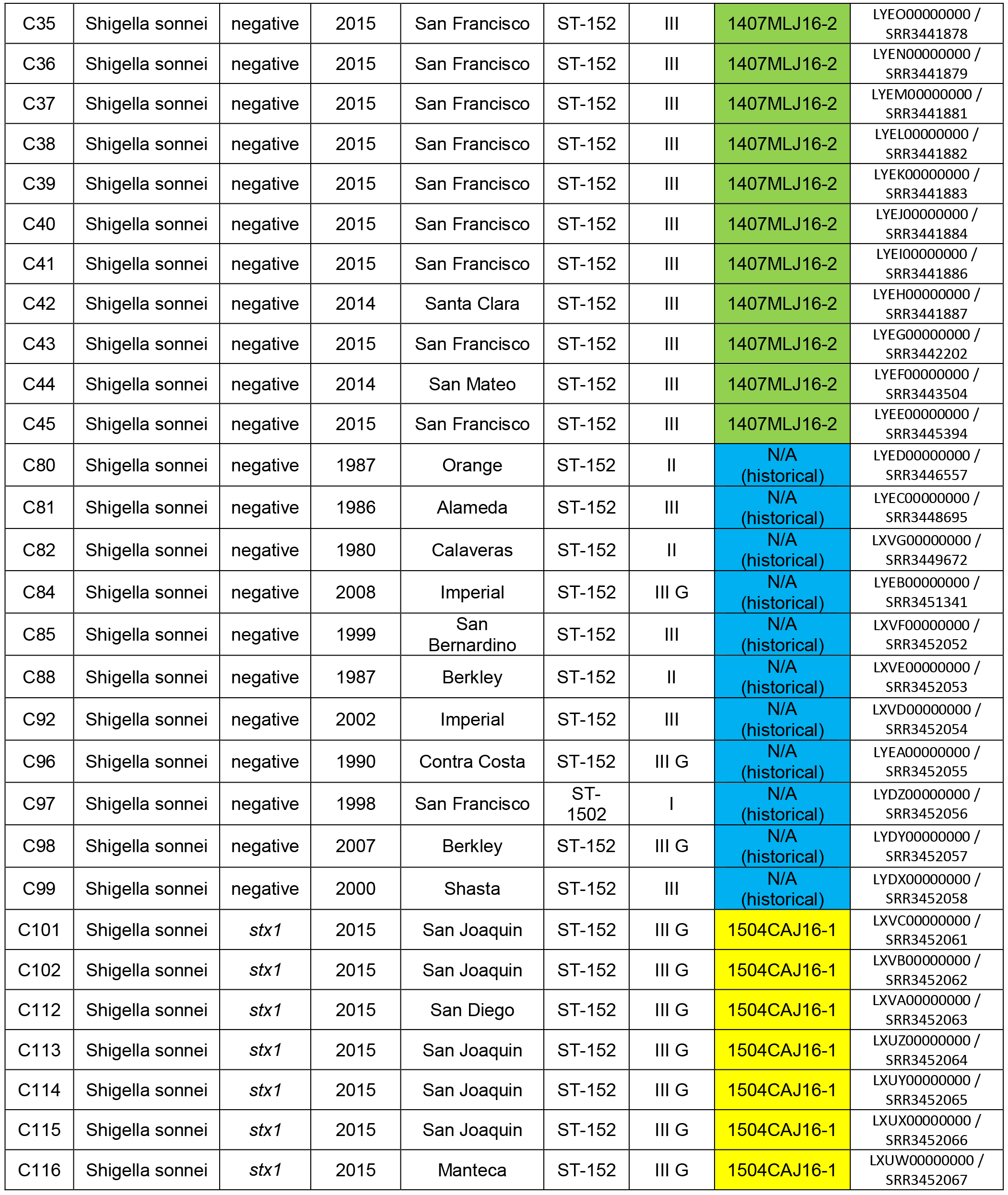

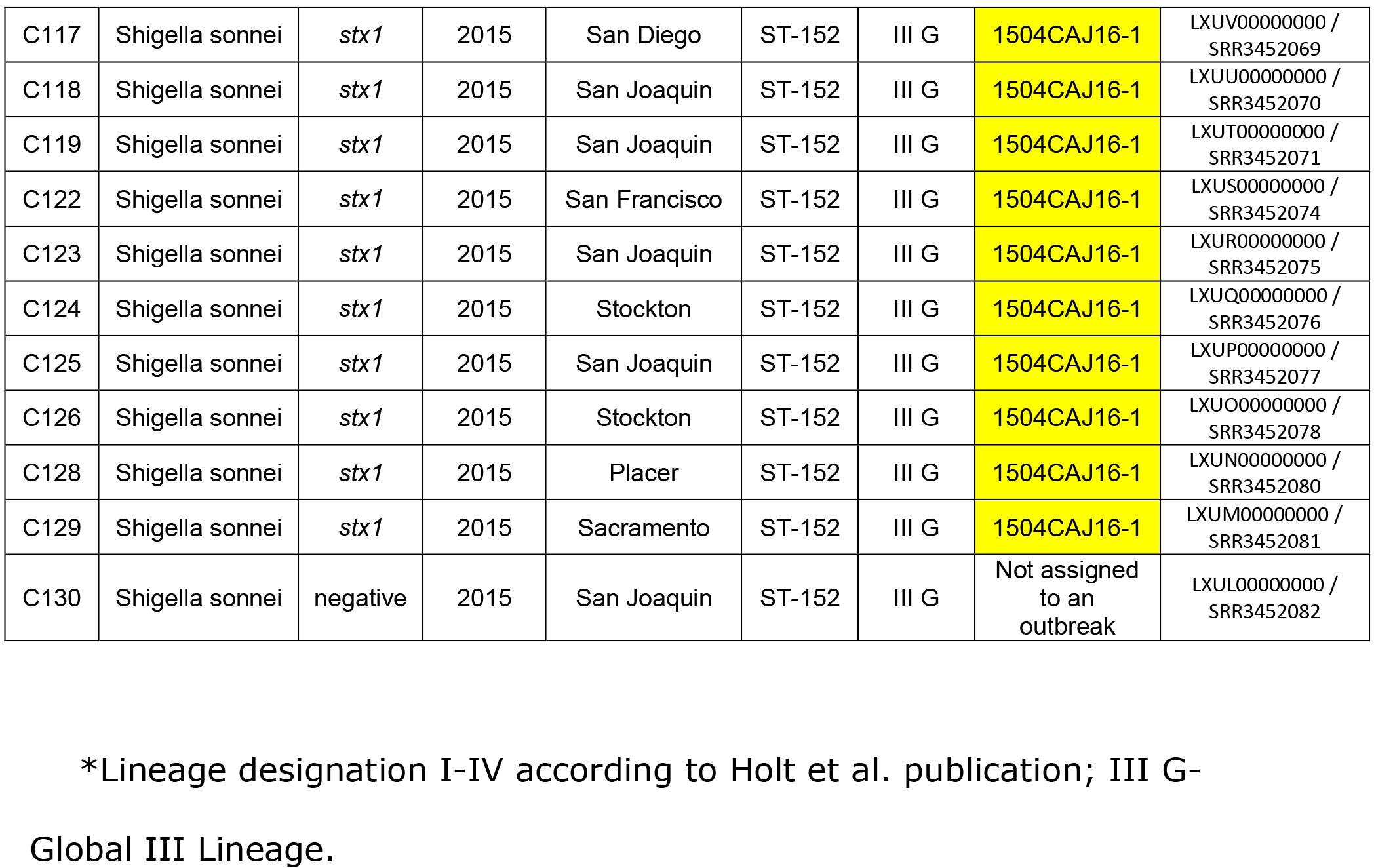

**Supplementary Table 2.**
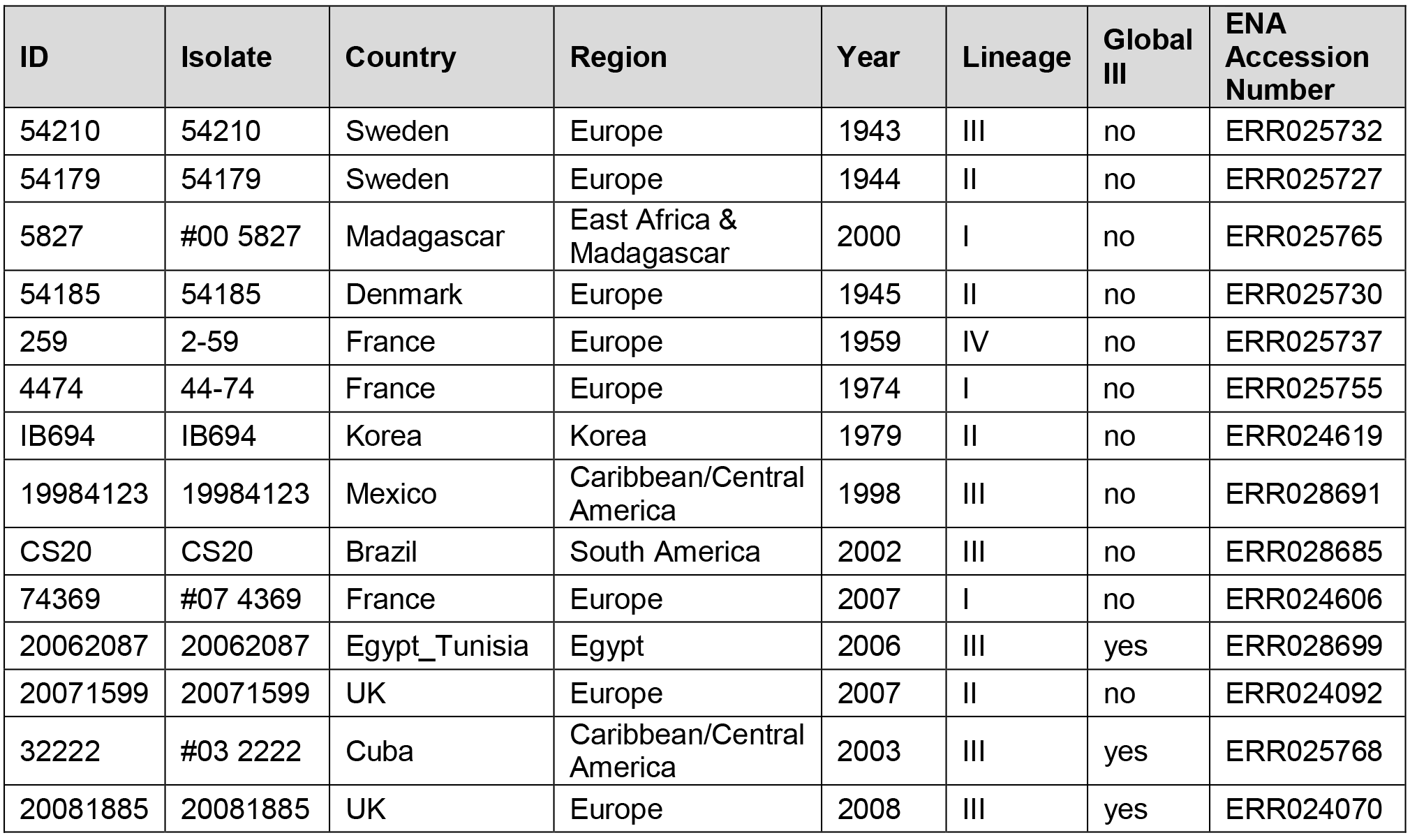
List of isolates from Holt et al. publication used for phylogenetic comparison.

**Supplementary Table 3.** Genes of representative SD/SJ *S. sonnei* isolates with homologs in *E. coli* O104:H4, but not in other *S. sonnei* (except *S. sonnei* strain 1DT-1-) Green highlight-STX-bacteriophage genes; yellow-*bla*_TEM-1_ plasmid.

| Res ult | Gene Object ID | Locus Tag | Gene Name | Length | COG | Pfam | % ID |
| --- | --- | --- | --- | --- | --- | --- | --- |
| 1 | 2634646959 | Ga0082014_10014 | hypothetical protein | 2793 | - | - | 97.9 |
| 2 | 2634646960 | Ga0082014_10015 | hypothetical protein | 421 | - | - | 93.6 |
| 3 | 2634646961 | Ga0082014_10016 | Bacteriocin class II with double-glycine leader peptide | 83 | - | pfam10439 | 100 |
| 4 | 2634646962 | Ga0082014_10017 | hypothetical protein | 148 | - | - | 99.3 |
| 5 | 2634646964 | Ga0082014_10019 | hypothetical protein | 133 | - | - | 100 |
| 6 | 2634646967 | Ga0082014_100112 | hypothetical protein | 205 | - | - | 100 |
| 7 | 2634646969 | Ga0082014_100114 | protein of unknown function (DUF1983) | 422 | - | pfam09327 | 99.5 |
| 8 | 2634646970 | Ga0082014_100115 | hypothetical protein | 563 | - | - | 99 |
| 9 | 2634646971 | Ga0082014_100116 | Phage tail fibre adhesin Gp38 | 170 | - | pfam05268 | 100 |
| 10 | 2634646973 | Ga0082014_100118 | hypothetical protein | 216 | - | - | 99.5 |
| 11 | 2634646974 | Ga0082014_100119 | hypothetical protein | 187 | - | - | 100 |
| 12 | 2634646975 | Ga0082014_100120 | hypothetical protein | 153 | - | - | 100 |
| 13 | 2634646978 | Ga0082014_100123 | hypothetical protein | 335 | - | - | 99.4 |
| 14 | 2634646979 | Ga0082014_100124 | hypothetical protein | 714 | - | - | 100 |
| 15 | 2634646981 | Ga0082014_100126 | hypothetical protein | 268 | - | - | 100 |
| 16 | 2634646985 | Ga0082014_100130 | hypothetical protein | 44 | - | - | 100 |
| 17 | 2634646987 | Ga0082014_100132 | lysozyme | 177 | COG3772 | pfam00959 | 96.6 |
| 18 | 2634646989 | Ga0082014_100134 | Protein of unknown function (DUF826) | 81 | - | pfam05696 | 96.3 |
| 19 | 2634646992 | Ga0082014_100137 | shiga toxin subunit B | 89 | - | pfam02258 | 58 |
| 20 | 2634646993 | Ga0082014_100138 | shiga toxin subunit A | 315 | - | pfam00161 | 54.8 |
| 21 | 2634646995 | Ga0082014_100140 | Phage NinH protein | 64 | - | pfam06322 | 100 |
| 22 | 2634646996 | Ga0082014_100141 | Bacteriophage Lambda NinG protein | 187 | - | pfam05766 | 100 |
| 23 | 2634646997 | Ga0082014_100142 | Protein of unknown function (DUF1367) | 149 | - | pfam07105 | 100 |
| 24 | 2634646999 | Ga0082014_100144 | ATP-dependent helicase IRC3 | 506 | COG1061 | pfam00271<br>pfam04851 | 98.2 |
| 25 | 2634647000 | Ga0082014_100145 | hypothetical protein | 125 | - | - | 100 |
| 26 | 2634647001 | Ga0082014_100146 | Cro protein | 67 | - | pfam09048 | 100 |
| 27 | 2634647002 | Ga0082014_100147 | SOS-response transcriptional repressor LexA (RecA-mediated autopeptidase) | 237 | COG1974 | pfam00717<br>pfam01381 | 100 |
| 28 | 2634647004 | Ga0082014_100149 | BsuBI/PstI restriction endonuclease C-terminus | 317 | - | pfam06616 | 100 |
| 29 | 2634647008 | Ga0082014_100154 | hypothetical protein | 70 | - | - | 100 |
| 30 | 2634647009 | Ga0082014_100155 | hypothetical protein | 73 | - | - | 100 |
| 31 | 2634647010 | Ga0082014_100156 | hypothetical protein | 93 | - | - | 100 |
| 32 | 2634647011 | Ga0082014_100157 | recombination protein RecT | 316 | COG3723 | pfam03837 | 99.1 |
| 33 | 2634647012 | Ga0082014_100158 | YqaJ-like recombinase domain-containing protein | 229 | - | pfam09588 | 100 |
| 34 | 2634647013 | Ga0082014_100159 | hypothetical protein | 195 | - | - | 99 |
| 35 | 2634647016 | Ga0082014_100162 | hypothetical protein | 147 | - | - | 90.1 |
| 36 | 2634647020 | Ga0082014_100166 | Ead/Ea22-like protein | 233 | - | pfam13935 | 95.5 |
| 37 | 2634647021 | Ga0082014_100167 | hypothetical protein | 171 | - | - | 100 |
| 38 | 2634647025 | Ga0082014_100171 | hypothetical protein | 143 | - | - | 100 |
| 39 | 2634647026 | Ga0082014_100172 | N6-adenosine-specific RNA methylase IME4 | 208 | COG4725 | pfam05063 | 100 |
| 40 | 2634647028 | Ga0082014_100174 | Protein of unknown function (DUF1627) | 129 | - | pfam07789 | 96.9 |
| 41 | 2634647030 | Ga0082014_100176 | protein of unknown function (DUF4222) | 83 | - | pfam13973 | 98.8 |

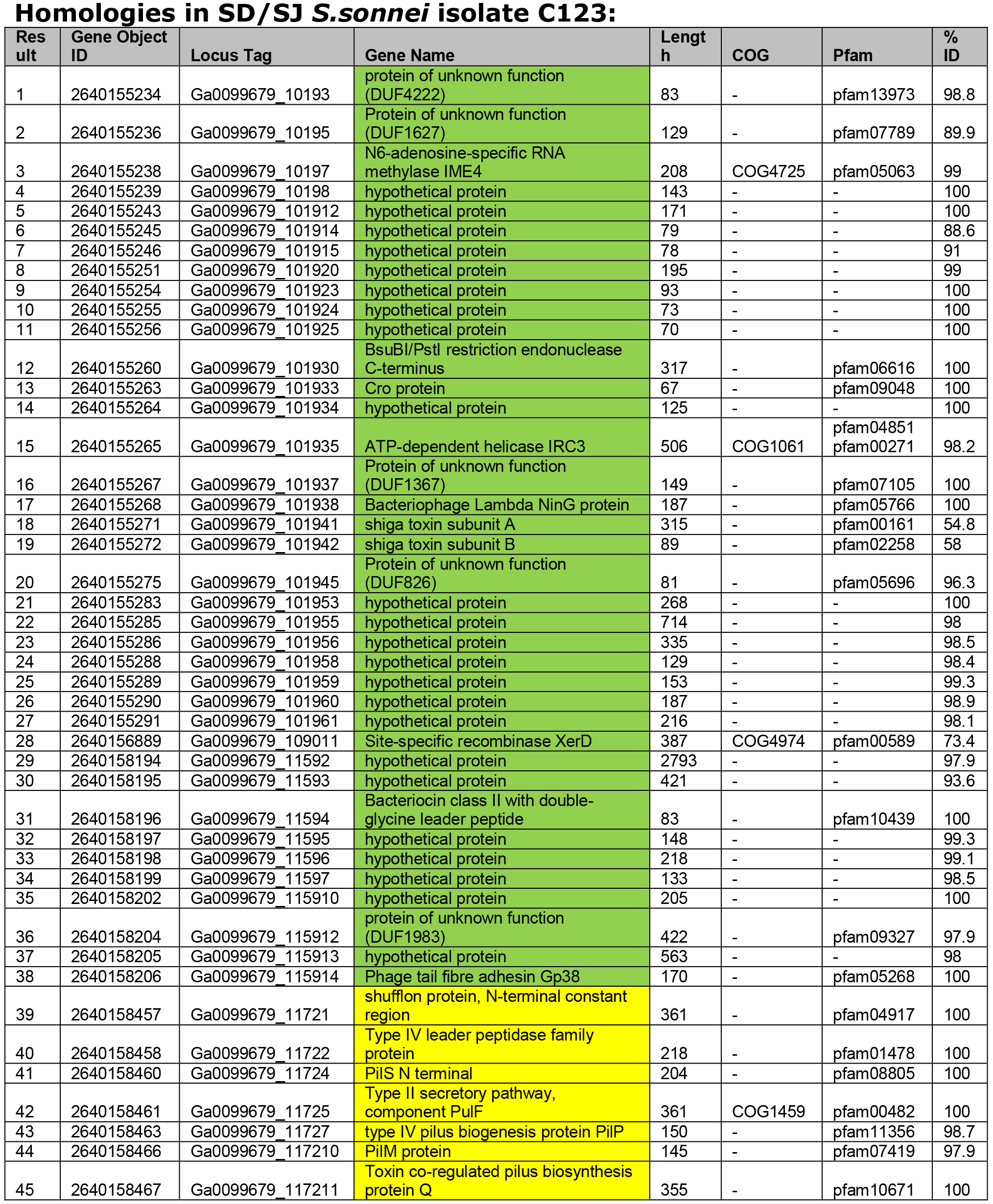

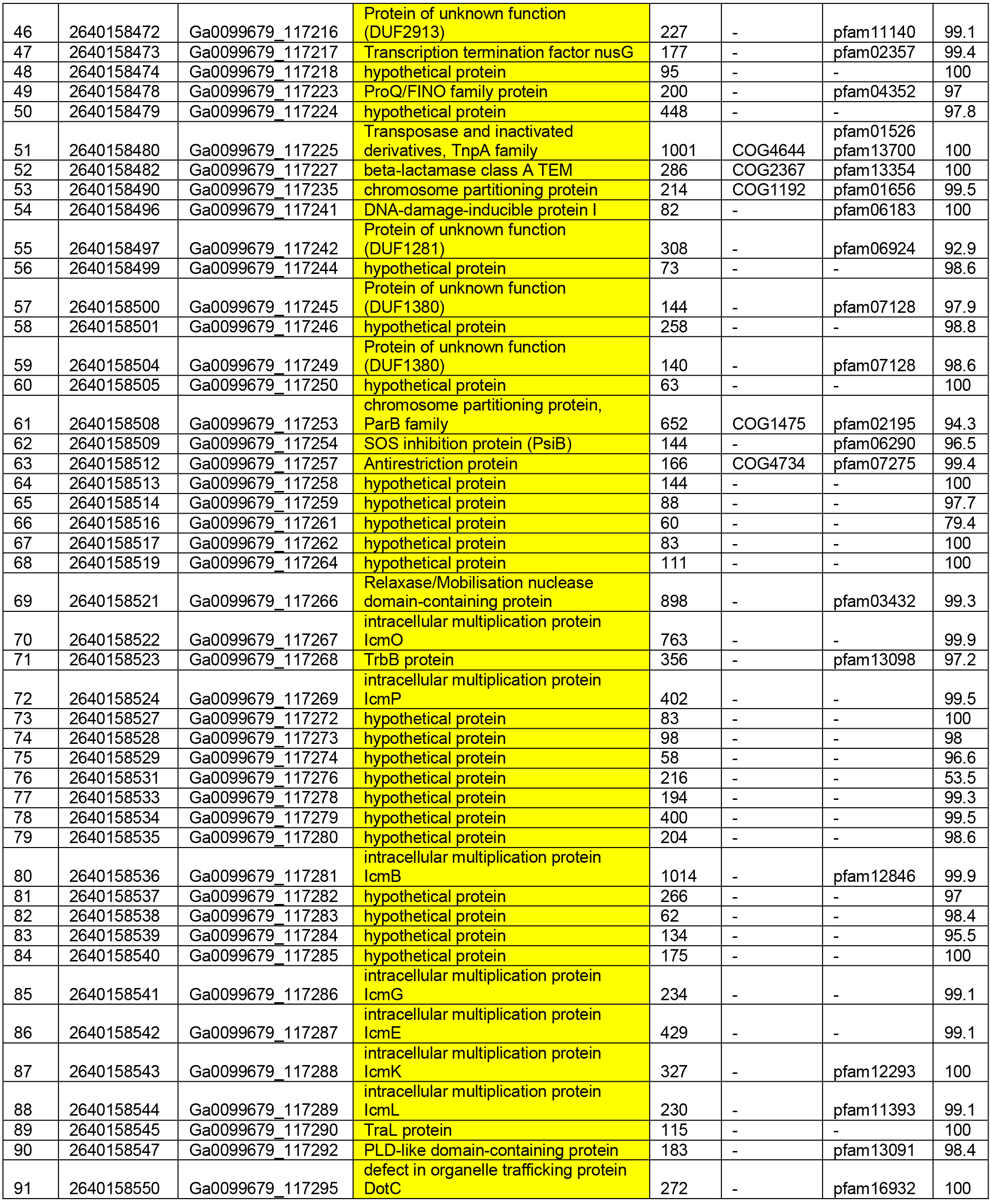

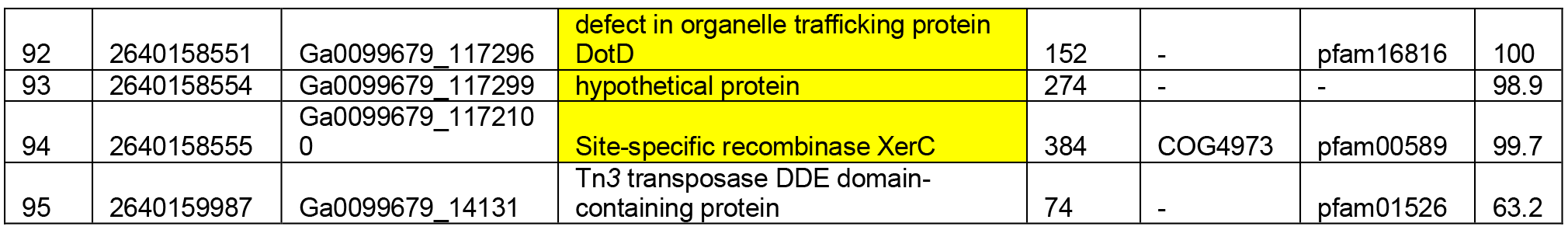

**Supplementary Table 4.**
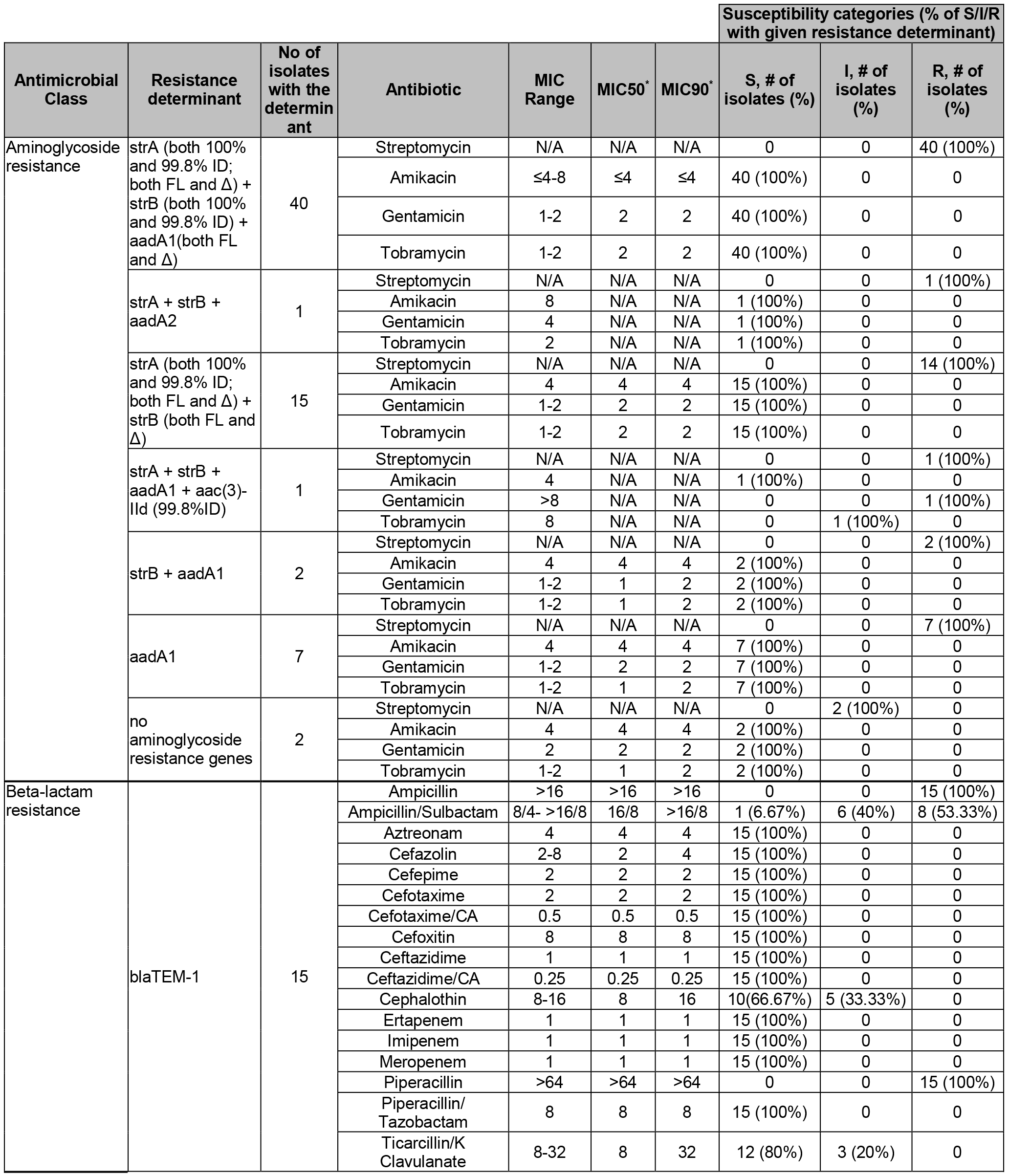
Antibiotic resistance genotype and phenotype of CA S. sonnei isolates.

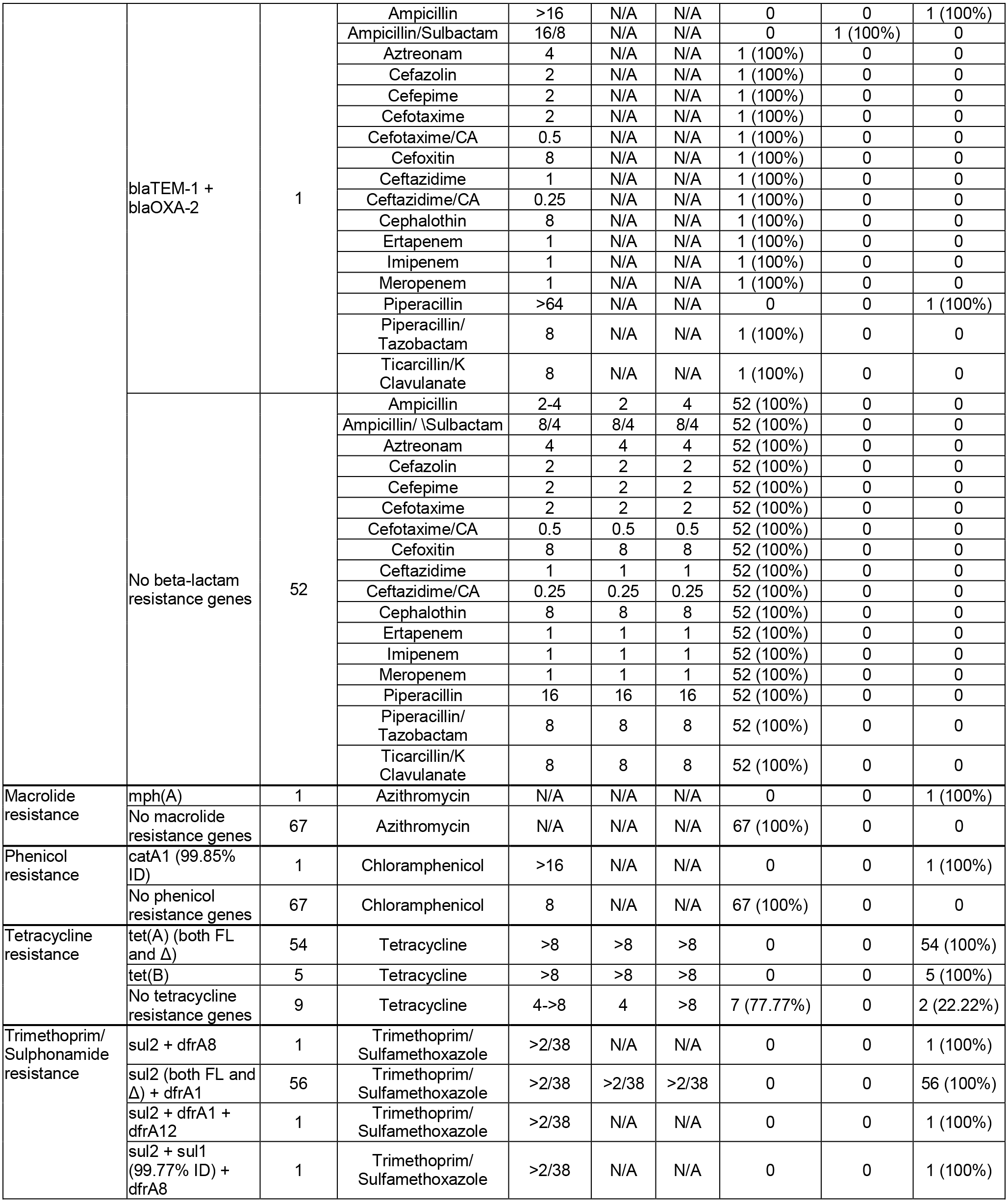

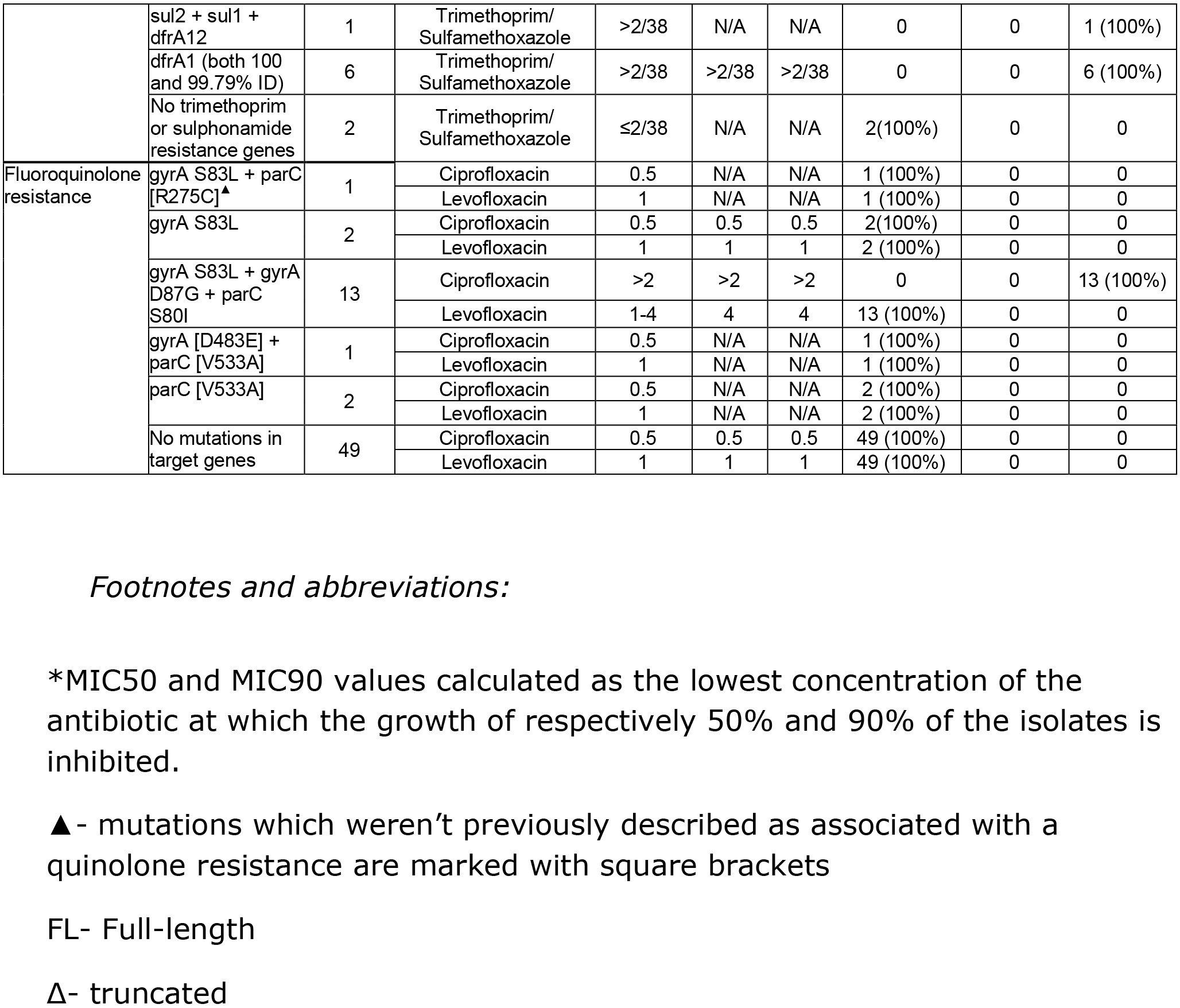

